# Divergence amid recurring gene flow: complex demographic histories for two North American pines (*Pinus pungens* and *P. rigida*) fit growing expectations among forest trees

**DOI:** 10.1101/2022.02.12.480138

**Authors:** Constance E. Bolte, Trevor M. Faske, Christopher J. Friedline, Andrew J. Eckert

**Affiliations:** Integrative Life Sciences Doctoral Program, Virginia Commonwealth University, Richmond, VA, USA; Ecology, Evolution & Conservation Biology Doctoral Program, University of Nevada, Reno, NV, USA; Department of Biology, Virginia Commonwealth University, Richmond, VA, USA

**Keywords:** conifer speciation, *Pinus pungens*, *Pinus rigida*, reproductive isolation, population genetics, species distributions

## Abstract

Long-lived species of trees, especially conifers, often display weak patterns of reproductive isolation, but clear patterns of local adaptation and phenotypic divergence. Discovering the evolutionary history of these patterns is paramount to a generalized understanding of speciation for long-lived plants. We focus on two closely related yet phenotypically divergent pine species, *Pinus pungens* and *P. rigida*, that co-exist along high elevation ridgelines of the southern Appalachian Mountains. In this study, we performed historical species distribution modeling (SDM) to form hypotheses related to population size change and gene flow to be tested in a demographic inference framework. We further sought to identify drivers of divergence by associating climate and geographic variables with genetic structure within and across species boundaries. Population structure within each species was absent based on genome-wide RADseq data. Signals of admixture were present range-wide, however, and species-level genetic differences associated with precipitation seasonality and elevation. When combined with information from contemporary and historical species distribution models, these patterns are consistent with a complex evolutionary history of speciation influenced by Quaternary climate. This was confirmed using inferences based on the multidimensional site-frequency spectrum, where demographic modeling inferred recurring gene flow since divergence (2.74 million years ago) and population size reductions that occurred during the last glacial period (∼35.2 thousand years ago). This suggests that phenotypic and genomic divergence, including the evolution of divergent phenological schedules leading to partial reproductive isolation, as previously documented for these two species, can happen rapidly, even between long-lived species of pines.

## Introduction

The process of speciation has been characterized as a continuum of divergence underpinned with the expectation that reproductive isolation strengthens over time leading to increased genomic conflict between species (Seehausen et al. 2014). While the term continuum suggests linear directionality, it is better thought of as a multivariate trajectory that is nonlinear, allowing stalls and even breakdown of reproductive barriers in the overall progression toward complete reproductive isolation (Cannon and Petit 2020; Kulmuni et al. 2020). Indeed, speciation can occur with or without ongoing gene flow and demographic processes such as expansions, contractions, isolation, and introgression leave detectable genetic patterns within and among populations of species that affect the evolution of reproductive isolation (Nosil 2012; e.g., Gao et al. 2012). Divergence histories with gene flow are an emerging pattern for species of forest trees with reproductive isolation often developing through prezygotic isolating mechanisms and reinforced by environmental adaptation (Abbott 2017; Cavender-Bares 2019). Together, these two processes can facilitate the development of genomic incompatibilities over time (Baack et al. 2015).

Climate and geography are well-established drivers of demographic processes and patterns (Hewitt 2001). For the past 2.6 million years, Quaternary climate has oscillated between glacial and interglacial periods causing changes in species distributions, but the significance of these changes and their influence on population differentiation has varied by region and taxon (Hewitt 2004; Lascoux et al. 2004). In North America, the effects of Quaternary climate on tree species distributions and patterns of genetic diversity have been profound but more drastic for species native to northern (i.e., previously glaciated) and eastern regions. For instance, the geographical distribution of white oak (*Quercus alba* L.), a native tree species to eastern North America, experienced greater shifts since the last interglacial period (LIG), approximately 120 thousand years ago (kya), compared to the distributional shifts of valley oak (*Quercus lobata* Née) in California (Gugger et al. 2013). For the latter, distributional, and hence niche, stability was correlated with higher levels of genetic diversity.

Given the climate instability of eastern North America since the LIG, a host of phylogeographic studies have reported genetic diversity estimates for taxa of this region, as well as the genetic structuring of populations due to geographic barriers such as the Appalachian Mountains and Mississippi River (Soltis et al. 2006) and postglacial expansion (e.g., Gougherty et al. 2020). The vast majority of tree taxa in these studies, however, were angiosperms, with the divergence history of only one closely related pair of conifer species native to this region, *Picea mariana* (Mill.) Britton, Sterns, & Poggenb. and *P. rubens* Sarg., being fully characterized (Perron et al. 2000; Lafontaine et al. 2015). The relative differences in geographical distributions and genetic diversities across *P. mariana* and *P. rubens,* as well as models of demographic inference, suggest a progenitor-derivative species relationship that initiated approximately 110 kya through population contractions and geographical isolation. Despite this history, these two species actively hybridize today. In general, speciation among conifer lineages remains an enigmatic process (Bolte and Eckert 2020), largely because there is a mismatch between species-level taxonomy and the existence of reproductive isolation, so that hybridization among species is common both naturally as well as artificially (Critchfield 1986). The ability to hybridize, moreover, is idiosyncratic, with examples ranging from well-developed incompatibilities among populations within species (e.g., *P. muricata* D. Don; Critchfield 1967) to the almost complete lack of incompatibilities among diverged and geographically distant species (*P. wallichiana* A. B. Jacks. from central Asia and *P. monticola* Douglas ex D. Don from western North America; Wright 1959). Thus, the tempo and mode for the evolution of reproductive isolation for conifers remains largely unexplained despite decades of research into patterns of natural hybridization, crossing rates, and the mechanisms behind documented incompatibilities (McWilliam 1959; Kriebel 1972; Hagman 1975; Critchfield 1986; Vasilyeva and Goroshkevich 2018).

The key to understanding the evolution of reproductive isolation, and hence a more developed explanation of the process of speciation for conifers, is the role of demography and gene flow during the divergence among lineages. Analytical approaches have been developed to infer past demographic processes from population genomic data, which can now easily be generated even for conifers (Parchman et al. 2018). While many studies have used demographic inference to describe the phylogeographic history of a single species (e.g., Gugger et al. 2013; Li et al. 2013; Bagley et al. 2020; Ju et al. 2019; Park and Donoghue 2019; Capblancq et al. 2020; Yang et al. 2020; Labiszak et al. 2021), some of these established methods have also been used to infer divergence histories between two or three species (e.g., Zou et al. 2013; Christe et al. 2017; Kim et al. 2018; Menon et al. 2018). Single species inferences have found that the last glacial maximum (LGM; ∼21 kya) affected distributional shifts and intraspecific gene flow dynamics, while multispecies studies have focused almost solely on how these climatic oscillations drove periods of increased and decreased interspecific gene flow which contributed to the formation of environmentally dependent hybrid zones, ancient and periodical introgression, or adaptive divergence in the development of reproductive isolation.

The number of potential divergence histories underlying even a modest number of species is vast. The preemptive formation of a hypothesis from historical species distribution modeling (SDM), however, can aid in defining a more realistic set of models from which to make inference, as well as to examine the impact of climate change on genetic diversity and demographic processes (Carstens and Richards, 2007). For example, Lima et al. (2017) modeled distributional changes for *Eugenia dysenterica* DC. between the LGM and today leading to a hypothesis that range stability was more likely than range expansion or contraction. Their SDM-informed hypothesis was supported by range-wide genetic data. Likewise, SDMs across several time points allow for estimation of habitat suitability change (i.e., a proxy for contraction or expansion) and distributional overlap of multiple species (i.e., potential gene flow). With these quantified changes, testable hypotheses can often emerge, leading to more focused investigations of speciation through justified parameter selection (Richards et al. 2007). Of course, there are inherent limitations associated with SDMs and interpreting historical distributions should be done cautiously but using SDMs to complement demographic inference is now common in the field of phylogeography (Hickerson et al. 2010; Gavin et al. 2014; Peterson and Anamza 2015). For example, where a species occurs is determined to some degree by its traits and thus at least partially its genetics, so that non-optimal inference can occur by ignoring putative adaptation within lineages during SDM formation and testing. Indeed, Ikeda et al. (2017) found that SDM predictions under future climate scenarios improved with acknowledgement of local adaptation in *Populus fremontii* S. Watson (i.e., three identified genetic clusters across the full species distributional range were modeled independently).

Here, we focus on two closely related, yet phenotypically diverged, pine species, Table Mountain pine (*Pinus pungens* Lamb.) and pitch pine (*Pinus rigida* Mill.). Recent estimates from multiple, time-calibrated phylogenies across nuclear and plastid DNA have placed the time of divergence in the range of 1.5 to 17.4 million years ago (mya; Hernandez-Leon et al. 2013; Saladin et al. 2017; Gernandt et al. 2018; Jin et al. 2021), with these studies either placing them as sister species (e.g., Hernandez-Leon et al. 2013; Saladin et al. 2017) or as part of a clade with *P. serotina* Michx. as sister to *P. rigida* (e.g., Gernandt et al. 2018; Jin et al. 2021). Changes in climate, fire regime, and geographic distributions have likely influenced species divergence (Keeley 2012). This is plausible given that *P. pungens* populations are restricted to high elevations of the Appalachian Mountains, while the much larger distribution of *P. rigida* ranges from Georgia into portions of eastern Canada. It is particularly interesting that these recently diverged species are found in sympatry, yet hybridization has rarely been observed in the field (Zobel 1969), although they can be reciprocally crossed to yield viable offspring (Critchfield 1963). An ecological study of three sympatric *P. pungens* and *P. rigida* populations indicated that the timing of pollen release was separated by approximately four weeks, enough to sustain partial reproductive isolation at these sites (Zobel 1969), which is a common contributor to prezygotic isolation among conifer species (Dorman and Barber 1956; Critchfield 1963). It was also noted that while *P. pungens* was most densely populated on arid, rocky, steep southwestern slopes, *P. rigida* was less confined to these areas (Zobel 1969), thus suggesting environmental adaptation through ecological character displacement may also be important in the divergence of these two closely related species.

Considering the dynamic interplay of climate, topography, and ecology potentially involved in the divergence of these two pine species, we asked three questions: 1) Which demographic processes were involved in the divergence of *P. pungens* and *P. rigida*? 2) Does the timing of demographic events align with shifts in climate? 3) To what extent are climatic and geographic variables associated with genetic differentiation? To answer these three questions, we hypothesized that *P. pungens* and *P. rigida* experienced divergence with gene flow followed by population contraction and isolation (i.e., different refugia) initiated during the LGM as an explanation for strongly diverged traits and phenological schedules. From historical SDM predictions across four time points since the LIG, we formed additional hypotheses to be tested within a demographic inference framework. Three hypotheses corresponded to SDM predictions from specific general circulation models (GCMs) and were compared to a fourth hypothesis formed from ensembled SDM predictions. We then used a multidimensional, folded site frequency spectrum from 2168 genome-wide, unlinked single nucleotide polymorphisms (SNPs) across 300 trees to infer demographic processes and timing of divergence. Our best-fit demographic model inferred initial divergence at 2.74 mya, aligning with the start of the Quaternary Period, and described divergence as occurring with ongoing gene flow and drastic population size reductions during the last glacial period (∼35.2 kya). SDM hypotheses were partially supported, especially for ongoing gene flow and population size reductions during the LGM. We conclude that climatic oscillations, differential adaptation to seasonality, and gene flow influenced the divergence of *P. pungens* and *P. rigida* and present evidence from SDM, genetic association analyses, and demographic inference as support.

## Methods

### Sampling

Range-wide samples of needle tissue were obtained from 14 populations of *Pinus pungens* and 19 populations of *P. rigida* (Fig. 1). From each population 4-12 trees were sampled, with each sampled tree distanced by approximately 50 m from the next to avoid potential kinship (Table 1). Needle tissue was dried using silica beads, then approximately 10 mg of tissue was cut and lysed for DNA extraction.

**Fig. 1.**
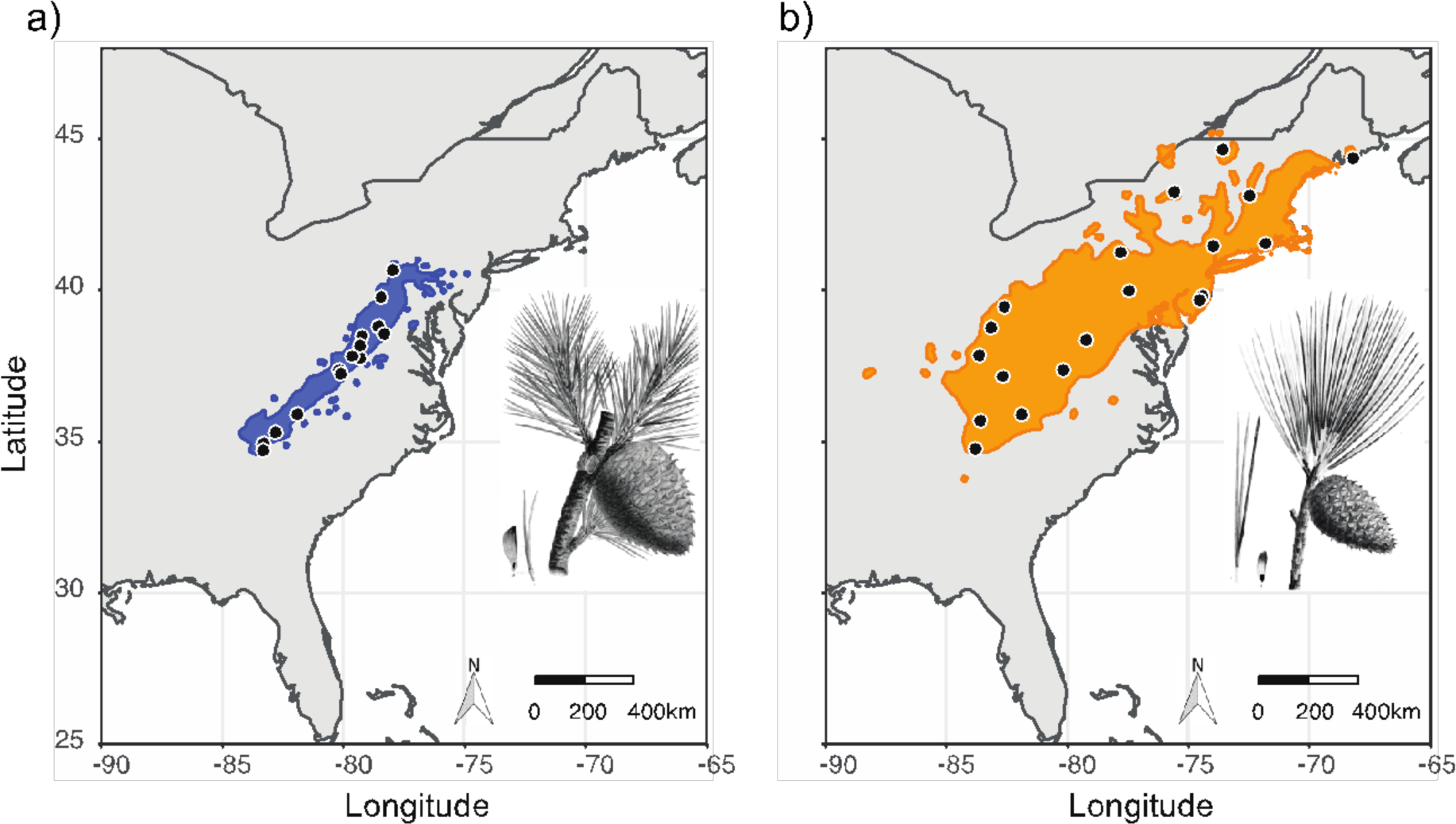
Known geographical distribution of focal species, a) *Pinus pungens* and b) *P. rigida*, (Little 1971) in relation to populations sampled (black dots) for genetic analysis; Phenotypic characterization of each species was illustrated by Pierre-Joseph Redouté (Michaux 1819).

**Table 1.** Location of sampled populations, number of trees (n) that were sampled, and the observed heterozygosity (Ho) versus the expected heterozygosity (He = 2pq) for Pinus pungens and P. rigida populations.

| Species | Code | Location | Lat | Long | $n$ | $H_o$ | $H_e$ |
| --- | --- | --- | --- | --- | --- | --- | --- |
| *P. pungens* | PU_BB | Briery Branch, VA | 38.48 | -79.22 | 8 | 0.110 | 0.108 |
| *P. pungens* | PU_BN | Buchanan State Forest, PA | 39.77 | -78.43 | 6 | 0.141 | 0.121 |
| *P. pungens* | PU_BV | Buena Vista, VA | 37.76 | -79.29 | 11 | 0.124 | 0.120 |
| *P. pungens* | PU_DT | Dragon's Tooth, VA | 37.37 | -80.16 | 7 | 0.101 | 0.098 |
| *P. pungens* | PU_EG | Edinburg Gap, VA | 38.79 | -78.53 | 8 | 0.139 | 0.124 |
| *P. pungens* | PU_EK | Elliott Knob, VA | 38.17 | -79.30 | 10 | 0.131 | 0.123 |
| *P. pungens* | PU_GA | Walnut Fork, GA | 34.92 | -83.28 | 10 | 0.129 | 0.123 |
| *P. pungens* | PU_LG | Looking Glass Rock, NC | 35.30 | -82.79 | 8 | 0.130 | 0.119 |
| *P. pungens* | PU_NM | North Mountain, VA | 37.82 | -79.63 | 12 | 0.130 | 0.121 |
| *P. pungens* | PU_PM | Poor Mountain, VA | 37.23 | -80.09 | 11 | 0.130 | 0.125 |
| *P. pungens* | PU_SC | Pine Mountain, VA | 34.70 | -83.30 | 8 | 0.128 | 0.122 |
| *P. pungens* | PU_SH | Shenandoah NP, VA | 38.55 | -78.31 | 5 | 0.160 | 0.128 |
| *P. pungens* | PU_SV | Stone Valley Forest, PA | 40.66 | -77.95 | 9 | 0.110 | 0.110 |
| *P. pungens* | PU_TR | Table Rock Mountain, NC | 35.89 | -81.88 | 12 | 0.113 | 0.114 |
| *P. rigida* | RI_BR | Bass River State Forest, NJ | 39.80 | -74.41 | 9 | 0.101 | 0.105 |
| *P. rigida* | RI_CT | Pachaug State Forest, CT | 41.54 | -71.81 | 10 | 0.096 | 0.107 |
| *P. rigida* | RI_DT | Dragon's Tooth, VA | 37.37 | -80.16 | 10 | 0.109 | 0.106 |
| *P. rigida* | RI_GA | Chattahoochee NF, GA | 34.75 | -83.78 | 9 | 0.096 | 0.103 |
| *P. rigida* | RI_GW | George Washington NF, VA | 38.36 | -79.20 | 10 | 0.102 | 0.103 |
| *P. rigida* | RI_HH | Hudson Highlands State Park, NY | 41.44 | -73.97 | 7 | 0.102 | 0.101 |
| *P. rigida* | RI_JF | Jefferson NF, VA | 37.15 | -82.64 | 10 | 0.095 | 0.100 |
| *P. rigida* | RI_KY | Daniel Boone NF, KY | 37.84 | -83.62 | 9 | 0.113 | 0.110 |
| *P. rigida* | RI_ME | Acadia NP, ME | 44.36 | -68.19 | 10 | 0.107 | 0.106 |
| *P. rigida* | RI_MI | Michaux State Forest, PA | 39.98 | -77.44 | 10 | 0.123 | 0.114 |
| *P. rigida* | RI_NJ | Wharton State Forest, NJ | 39.68 | -74.53 | 9 | 0.098 | 0.101 |
| *P. rigida* | RI_NY | Macomb State Park, NY | 44.63 | -73.58 | 9 | 0.101 | 0.104 |
| *P. rigida* | RI_OH | South Bloomingville, OH | 39.45 | -82.59 | 8 | 0.093 | 0.096 |
| *P. rigida* | RI_RS | Rome Sand Plains, NY | 43.23 | -75.56 | 9 | 0.097 | 0.103 |
| *P. rigida* | RI_SH | Shawnee State Park, OH | 38.75 | -83.13 | 9 | 0.082 | 0.094 |
| *P. rigida* | RI_SP | Sproul State Forest, PA | 41.24 | -77.78 | 9 | 0.106 | 0.105 |
| *P. rigida* | RI_TN | Great Smoky Mountains NP, TN | 35.68 | -83.58 | 8 | 0.099 | 0.104 |
| *P. rigida* | RI_TR | Table Rock Mountain, NC | 35.89 | -81.89 | 10 | 0.113 | 0.112 |
| *P. rigida* | RI_VT | Bellows Falls, VT | 43.11 | -72.44 | 10 | 0.098 | 0.104 |

### DNA sequence data

Genomic DNA was extracted from all 300 sampled trees using DNeasy Plant Kits (Qiagen) following the manufacturer’s protocol. Four ddRADseq libraries (Peterson et al. 2012), each containing up to 96 multiplexed samples, were prepared using the procedure from Parchman et al. (2012). EcoRI and MseI restriction enzymes were used to digest all four libraries before performing ligation of adaptors and barcodes. After PCR, agarose gel electrophoresis was used to separate then select DNA fragments between 300-500 bp in length. The pooled DNA was isolated using a QIAquick Gel Extraction Kit (Qiagen). Single-end sequencing was conducted on Illumina HiSeq 4000 platform by Novogene Corporation (Sacramento, CA). Raw fastq files were demultiplexed using GBSX (Herten et al. 2015) version 1.2, allowing two mismatches (-mb 2). The dDocent bioinformatics pipeline (Puritz et al. 2014) was subsequently used to generate a reference assembly and call variants. The reference assembly was optimized using shell scripts and documentation within dDocent (cutoffs: individual = 6, coverage = 6; clustering similarity: -c 0.92), utilizing cd-hit-est (Fu et al. 2012) for assembly. The initial variant calling produced 87,548 single nucleotide polymorphisms (SNPs) that were further filtered using *vcftools* (Danecek et al. 2011) version 0.1.15. We retained only biallelic SNPs with sequencing data for at least 50% of the samples, minor allele frequency (MAF) > 0.01, summed depth across samples > 100 and < 10000, and alternate allele call quality ≥ 50. Additionally, stringent filtering steps were taken to minimize the potential misassembly of paralogous genomic regions. Removing loci with excessive coverage and retaining only loci with two alleles present, as above, should ameliorate the influence of misassembled paralogous loci in our data (Hapke and Thiele 2016; McKinney et al. 2018). Lastly, we retained loci with *F*IS *>* –0.5, as misassembly to paralogous genomic regions can lead to abnormal levels of heterozygosity (Hohenlohe et al. 2013; McKinney et al. 2017). To account for linkage disequilibrium among the 20,932 SNPs that passed quality controls, which if not properly acknowledged can lead to erroneous inferences of demographic history (Gutenkunst et al. 2009), we thinned the dataset to one SNP per contig (--thin 100). The reduced 2168 SNP dataset was used in all analyses.

### Population structure and genetic diversity

Patterns of genetic diversity and structure within and between *P. pungens* and *P. rigida* were assessed using a suite of standard methods. Overall patterns of genetic structure were investigated using principal component analysis (PCA), as employed in the prcomp function of the *stats* version 4.0.4 package, on centered and scaled genotypes following Patterson et al. (2006) in R version 3.6.2 (R Development Core Team, 2021). Genetic diversity within each species was examined using multilocus estimates of observed and expected heterozygosity (*H*o and *H*e) for each population using a custom R script (www.github.com/boltece/Speciation_2pines). Individual-based assignment was conducted using *fastSTRUCTURE* (Raj et al. 2014), with cluster assignments ranging from *K* = 2 to *K* = 7. Ten replicate runs of each cluster assignment were conducted. The cluster assignment with the highest log-likelihood value was determined to be the best fit. Individual admixture assignments were then aligned and averaged across the 10 runs using the *pophelper* version 1.2.0 (Francis 2017) package in R. Third, multilocus, hierarchical fixation indices (*F*-statistics) were defined by nesting trees into populations and populations into species, with *F*CT describing differentiation between species and *F*SC describing population differentiation within species (Yang 1998). *F*-statistics and associated confidence intervals (95% CIs) from bootstrap resampling across SNPs (*n* = 100 replicates) were calculated in the *hierfstat* version 0.5-7 package (Goudet and Jombart 2020) in R.

To assess influences on within-species genetic structure, Mantel tests (Mantel 1967) were used to examine Isolation-by-Distance (IBD; Wright 1943) and Isolation-by-Environment (IBE; Wang and Bradburd 2014). In these analyses, the Mantel correlation coefficient (*r*) was calculated between linearized pairwise *F*ST, estimated with the method of Weir and Cockerham (1984) using the *hierfstat* package in R, and either geographical (IBD) or environmental (IBE) distances. For geographical distances, latitude, and longitude records for each tree in a population were averaged to obtain one representative coordinate per population. Geographic distances among populations were then calculated using the Vincenty (ellipsoid) method within the *geosphere* version 1.5- 10 package (Hijmans 2019) in R. Environmental distances were calculated as Euclidean distances using extracted raster values associated with the mean population coordinates from 19 bioclimatic variables, downloaded from WorldClim at 30 arc second resolution (version 2.1; Fick and Hijmans 2017). Values associated with the mean population coordinates for were extracted using the *raster* version 2.5-7 R package. Environmental data were centered and scaled prior to estimation of distances. Additionally, we used a Mantel test to assess correlation between population-based environmental distances and population-based geographic distances.

### Associations between genetic structure and environment

To further test the multivariate relationships among genotype, climate, and geography within and across species, redundancy analysis (RDA) was conducted using the *vegan* version 2.5-7 package (Oksanen et al. 2020) in R version 4.0.4 (R Core Development Team, 2021). Genotype data were coded as counts of the minor allele for each sample (i.e., 0,1, or 2 copies) and then standardized following Patterson et al. (2006). Climate raster data (i.e., 19 bioclimatic variables at 30 arc second resolutions), as well as elevational raster data from WorldClim, were extracted, as mentioned previously, from geographic coordinates for each sampled tree and then tested for correlation using Pearson’s correlation coefficient (*r*). Five bioclimatic variables that are known to influence diversification in the genus *Pinus* (Jin et al. 2021; Menon et al. 2018), but that were also not highly correlated (*r* < |0.75|), were retained for analysis: mean diurnal range (Bio2), maximum temperature of the warmest quarter (Bio10), and minimum temperature of the coldest quarter (Bio11), precipitation seasonality (Bio15), and precipitation of the driest quarter (Bio17). The full explanatory data set included these five bioclimatic variables, latitude, longitude, and elevation. The multivariate relationship between genetic variation, climate, and geography was then evaluated through RDA. Statistical significance (*α* = 0.05) of the RDA model, as well as each axis within the model, was assessed using permutation-based analysis of variance (ANOVA) with 999 permutations (Legendre and Legendre 2012). The influence of predictor variables, as well as their confounded effects, in RDA were quantified using variance partitioning as employed in the *varpart* function of *vegan* package in R.

### Species distribution modeling

To help formulate testable hypotheses during inference of demography from genomic data (see Richards et al. 2007), species distribution modeling (SDM) was performed for each species to identify areas of suitable habitat under current climate conditions and across three historical time periods (HOL, ∼6 kya, interglacial; LGM, ∼21 kya, glacial; and LIG, ∼120 kya, interglacial). These temporal inferences were then used to help identify plausible demographic responses. For example, if overlap in modeled habitat suitability changed over time, the hypothesis for demographic inference would include changes in gene flow parameters over time. If the amount of suitable habitat changed over time, the hypothesis would also include changes in effective population size to allow for potential expansions or contractions. This in effect helps to constrain the possible parameter space for exploration.

Occurrence records for *P. pungens* were downloaded from GBIF.org (18th December 2018; GBIF occurrence download, https://doi.org/10.15468/dl.urehu0) and combined with known occurrences published by Jetton et al. (2015). For *P. rigida*, all occurrence records were downloaded from GBIF.org (29th December 2015; GBIF occurrence download, http://doi.org/10.15468/dl.ak0weh). Records were examined for presence within or close to the known geographical range of each species (Little 1971). Records far outside the known geographic range were pruned. The remaining locations were then thinned to one occurrence per 10 km to reduce the effects of sampling bias using the *spThin* version 0.1.0.1 package (Aiello-Lammens et al. 2015) in R. The resulting occurrence dataset included 84 records for *P. pungens* and 252 records for *P. rigida* (Online Resource 2). All subsequent analyses were performed in R version 3.6.2 (R Development Core Team, 2021).

The same bioclimatic variables (Bio2, Bio10, Bio11, Bio15, Bio17) selected for RDA were used in species distribution modeling but were downloaded from WorldClim version 1.4 (Hijmans et al. 2005) at 2.5 arc minute resolution. The change in resolution from above was necessary because paleo-climate data in 30 arc second resolution were not available for the LGM. Paleoclimate raster data for the LGM (∼21 kya) and Holocene (HOL, ∼6 kya) were downloaded for three General Circulation Models (GCMs; CCSM4, MIROC-ESM, and MPI-ESM). Ensembles were built by averaging the habitat suitability predictions from the three GCMs for each time period (e.g., Menon et al. 2018). SDM predictions associated with each individual GCM, for both the HOL and LGM, were analyzed for incongruences as recommended in Varela et al. (2015). Paleoclimate data for the LIG (∼120 kya) were only available at 30 arc second resolution and required downscaling to 2.5 arc minute resolution using the aggregate function (fact = 5) of the *raster* package. Only one GCM is available for the LIG from WorldClim (NCAR-CCSM; Otto-Bliesner et al. 2006); therefore, no ensemble was built.

Raster layers were cropped to the same extent using the *raster* package to include the most northern and eastern extent of *P. rigida*, and the most western and southern extent of *P. pungens.* Species distribution models (SDMs) were built using MAXENT version 3.4.1 (Phillips et al. 2017) and all possible features and parameter combinations were evaluated using the *ENMeval* version 2.0.0 R package (Kass et al. 2021). Metadata about model fitting and evaluation are available within Online Resource 2.

The selected features used in predictive modeling were those associated with the best-fit model as determined using AIC. Raw raster predictions were standardized to have the sum of all grid cells equal the value of one using the *raster.standardize* function in the *ENMTools* version 1.0.5 (Warren et al. 2021) R package. Standardized predictions were then transformed to a cumulative raster prediction with habitat suitability scaled from 0 to 1, allowing for quantitative SDM comparisons across species and time. Next, SDM cumulative raster predictions were converted into coordinate points using the *sf* version 0.9-7 R package to calculate the number of points with habitat suitability values greater than 0.5 (i.e., moderate to high suitability areas). Population size expansion or contraction was hypothesized if the number of points increased or decreased over time, respectively. Overlap (i.e., shared points across species) in SDM predictions for each time period was measured using the *inner_join* function in the *dplyr* version 1.0.5 R package. The extent of modeled species distributional overlap was also quantified using the *raster.overlap* function in *ENMTools,* thus providing measures for calculation of Schoener’s *D* (1968) and Warren’s *I* (Warren et al. 2008). Four testable hypotheses were formed from these quantifications. Three of which were formed from predictions associated with each GCM used in HOL and LGM SDMs. The fourth hypothesis was formed from ensembled SDM predictions for the HOL and LGM.

### Demographic modeling

Demographic modeling was conducted using Diffusion Approximation for Demographic Inference (*δαδi* v.2.0.5; Gutenkunst et al. 2009). A model of pure divergence (SI; strict isolation) was compared against twelve other demographic models representing different potential divergence scenarios with or without gene flow and effective population size changes (Online Resource 4, Fig. S1). Based on SDM predictions across four time points, we hypothesized that a model that allowed changes in effective population size and rate of gene flow before the LIG would best fit the genetic data. Ten replicate runs of each model were performed in *δαδi* with a 200 x 220 x 240 grid space and the nonlinear Broyden-Fletcher-Goldfarb-Shannon (BFGS) optimization routine. Model selection was conducted using Akaike information criterion (AIC; Akaike 1974). The best replicate run (highest log composite likelihood) for each model was then used to calculate ΔAIC (AICmodel i – AICbest model) scores (Burnham and Anderson 2002). From the best supported model, upper and lower 95% confidence intervals (CIs) for all parameters were obtained using the Fisher Information Matrix (FIM)-based uncertainty analysis. Unscaled parameter estimates and their 95% CIs were obtained using a per lineage substitution rate of 7.28 x 10^10^ substitutions/site/year rate for *Pinaceae* (De La Torre et al. 2017) and a generation time of 25 years (Ma et al. 2006). The mutation rate from De La Torre et al. (2017) was placed on a scale of generations using the number of years per generation (i.e., mutation rate per year multiplied by the number of years per generation). Genome length (L) a requirement for determining *Nref* (= θ/4μL) from *δαδi* parameters, was calculated as the sum across contigs (i.e., RADtags) of the number of bp per SNP. This quantity was calculated for each contig by dividing 92 bp (i.e., the trimmed length of eachcontig) by the number of SNPs in the contig from the unthinned SNP dataset (*n* = 20,932 SNPs in total). This was necessary because only a single SNP was retained per contig and counting all bp in a contig would upwardly bias the genome length (i.e., the SNPs were dropped but the bp they occupy would be counted).

## Results

### Population structure and genetic diversity

A clear separation at the species level was apparent along PC1, which explained 4.232% of the variation across the 2168 SNP x 300 tree data set (Fig. 2a). Of the 2168 SNPs analyzed, 380 of them were fixed for the same allele across all samples of *P. pungens*, and 196 SNPs were fixed (i.e., not polymorphic) across samples of *P. rigida*. The majority of biallelic SNPs fixed in one species were segregating the global minor allele at low frequency in the other species. The other 1592 SNPs were polymorphic in both species. Lack of population clustering within each species was observed when the points in PCA-space were labeled by population (Online Resource 4, Fig. S2). Using hierarchical *F*- statistics, the estimate of differentiation between species (*F*CT) was 0.117 (95% CI: 0.099 – 0.136), which was similar to that among all sampled populations (*F*ST = 0.123, 95% CI: 0.106 – 0.143), thus highlighting structure is largely due to differences between species. Differentiation among populations within species was consequently much lower (*F*SC = 0.007 (95% CI: 0.0055-0.0088) whether analyzed jointly (*F*SC) or separately (see Table 2). In the analysis of structure, *K* = 2 had the highest log-likelihood values (Fig. 2b).

**Fig. 2.**
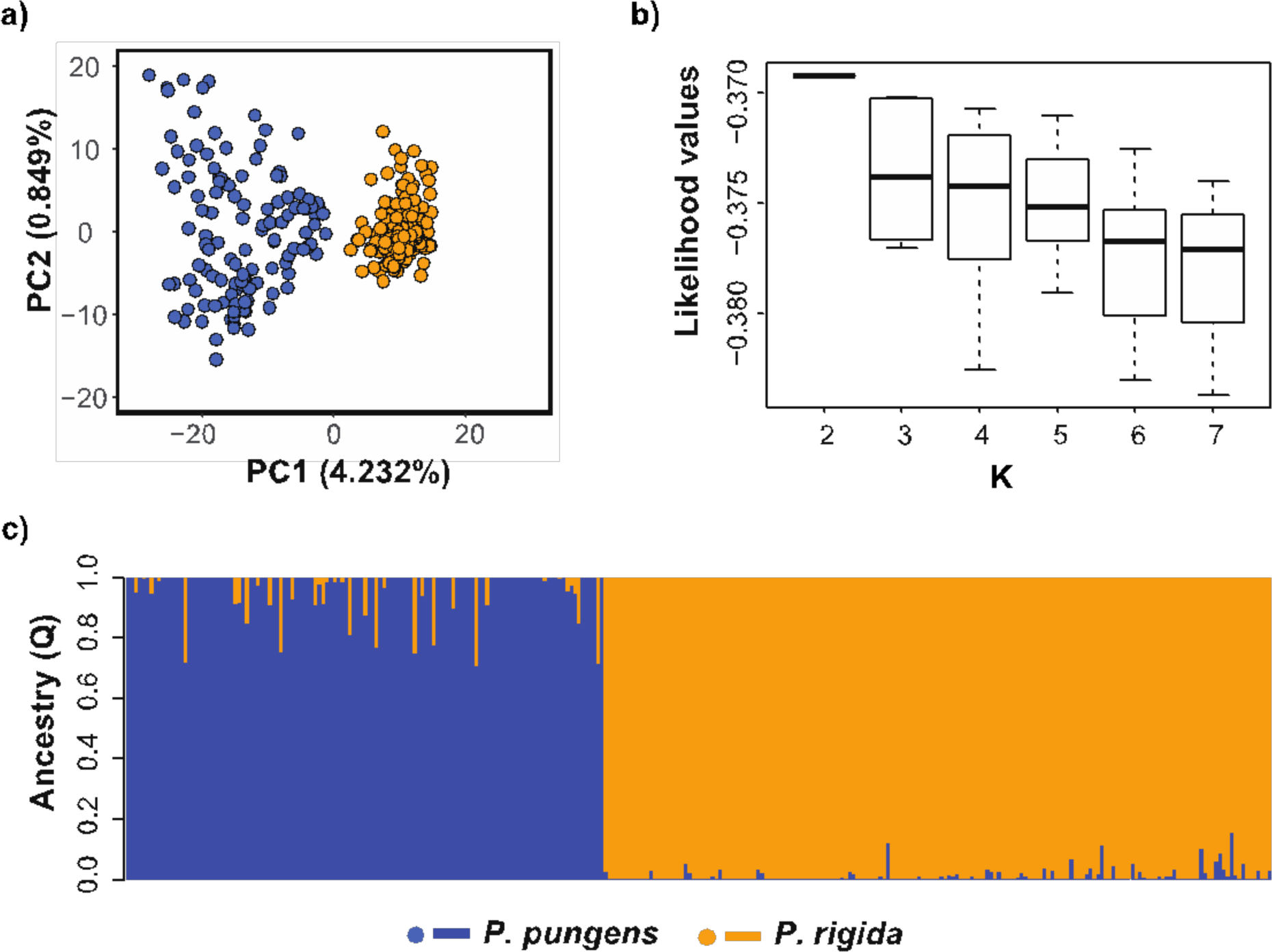
Measures of genetic differentiation and diversity among sampled trees of *P. pungens* and *P. rigida*: a) Principal components analysis of 2168 genome-wide single nucleotide polymorphism (SNPs) for *Pinus pungens* (blue, left side of PC1) and *P. rigida* (orange, right side of PC1); b) log-likelihood values across ten replicate runs in fastSTRUCTURE for *K* = 2 through *K* = 7; c) results of averaged *K* = 2 ancestry (*Q*) assignments for each sample arranged latitudinally in each species.

**Table 2.**
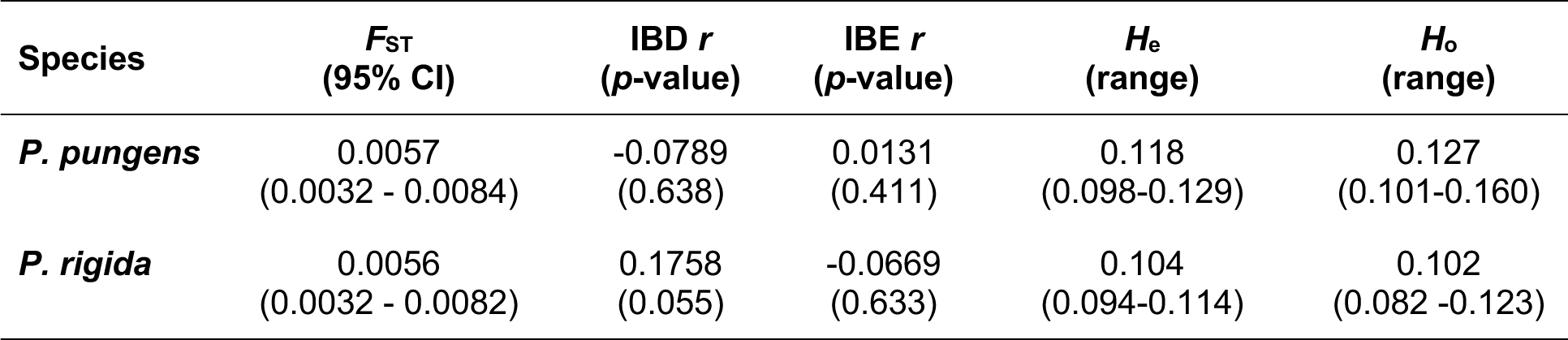
Summary statistics of genetic differentiation for the sampled populations of P. rigida and P. pungens. Expected (He) and observed heterozygosity (Ho) values are the averages across 2168 SNPs averaged across populations.

Admixture in small proportions (assigning to the other species by 2-10%) was observed in 41 out of the 300 samples (13.67% of samples) across both species. There were 16 trees with ancestry coefficients higher than 10% assignment to the other species: four *P. rigida* samples (2.29% of sampled *P. rigida*) and twelve *P. pungens* samples (9.60% of sampled *P. pungens*). Admixture proportions were moderately correlated to latitude (Pearson’s *r* = -0.414), longitude (Pearson’s *r* = -0.291), and elevation (Pearson’s *r* = 0.445). All three correlative relationships were significant (*p* < 0.001). Ancestry assignments for each tree at *K* = 3 through *K* = 7 are available in Online Resource 4 (Online Resource 4, Fig. S3). All cluster assignments analyzed did not reveal intraspecific population structure. To be certain the signals of admixture were not artifacts of missing data, we plotted the relationship of missing data to the ancestral coefficient for each tree. For the samples with admixture present, the assigned ancestral coefficients at *K* = 2 do not appear to be artifacts of missing data (Online Resource 4, Fig. S4). Admixture was present in trees with both low and moderate levels of missing data.

Pairwise *F*ST estimates for *P. pungens* ranged from 0 to 0.0457, while a similar but narrower range of values (0 – 0.0257) was noted for *P. rigida*. The highest pairwise *F*ST value across both species was between two *P. pungens* populations located in Virginia, PU_DT and PU_BB (Table 1). Interestingly, PU_DT in general had higher pairwise *F*ST values (0.0146 – 0.0457) compared to all the other sampled *P. pungens* populations (Online Resource 1). For *P. rigida*, the RI_SH population located in Ohio had higher pairwise *F*ST values for 16 out of the 18 comparisons (0.0123 – 0.0257). The two populations that had low pairwise *F*ST values with RI_SH were geographically proximal: RI_OH located in Ohio (pairwise *F*ST = 0, distance: 90.1 km) and RI_KY located in Kentucky (pairwise *F*ST = 0.0089, distance: 107.7 km). The highest pairwise *F*ST value among *P. rigida* populations was between RI_SH and RI_HH, which are geographically distant from one another. From the Mantel tests for IBD and IBE, Pearson correlations were low (Table 2). The correlation with geographical distances was highest for *P. rigida* (Mantel *r* = 0.176, *p* = 0.055). From the Mantel test, the correlation between geographic distance and environmental distance was high for both *P. rigida* (*r* = 0.611, *p* = 0.001) and *P. pungens* (*r* = 0.893, *p* = 0.001).

Heterozygosity estimates for each population are listed in Table 1 and were only moderately correlated with geography and elevation. Observed heterozygosity of *P. pungens* (*H*o = 0.127 ± 0.015 SD), averaged across SNPs and populations, was higher than the average expected heterozygosity (*H*e = 0.118 ± 0.008 SD), both of which were higher than the almost equal values for *P. rigida* (*H*o = 0.102 ± 0.009 SD; *H*e = 0.104 ± 0.005 SD; Table 2). Across both species, observed heterozygosity was mildly associated with geography and elevation. For *P. rigida*, the highest correlation was with elevation (*r* = 0.300, *p-*value = 0.212), followed by correlation with longitude (*r* = 0.113, *p-*value = 0.646). Observed heterozygosity in *P. pungens* had a negative correlative relationship with elevation (*r* = -0.105, *p*-value = 0.721) and positive correlative relationship with longitude (*r* = 0.175, *p*-values = 0.549). Correlations between latitude and heterozygosity were low in both species (*r* = -0.008 for *P. rigida*; *r* = 0.08 for *P. pungens*; *p-*values > 0.785).

### Associations between genetic structure and environment

The combined effects of climate and geography explained 1.52% (adj. *r*^2^) to 4.16% (*r*^2^) of the genetic variance across 2168 SNPs and 300 sampled trees. The first RDA axis accounted for the bulk of the explanatory variance (42.3%, Fig. 3) and was the only RDA axis with a *p*-value (*p* < 0.001) less than commonly accepted thresholds of significance (e.g., α = 0.05). The first RDA was dominated by effects of elevation and precipitation seasonality (Bio15). Average elevation associated with *P. pungens* samples was 724.68 m (± 224.17 SD), while average elevation across *P. rigida* samples was lower (399.69 m, ± 292.26 SD). The average for precipitation seasonality was 11.33 (± 1.83 SD) for *P. pungens*, and higher for *P. rigida* (14.23 ± 3.97 SD). Considering the standard deviations around the mean, overlap in values for elevation and precipitation seasonality provide some context to present day overlap in species distributions along the southern Appalachian Mountains. Comparisons of predictor loadings across both RDA axes show latitude, longitude, and mean temperature of the coldest quarter (Bio11) as also important to explaining the variance both within (RDA 2) and across species (RDA 1).

**Fig. 3.**
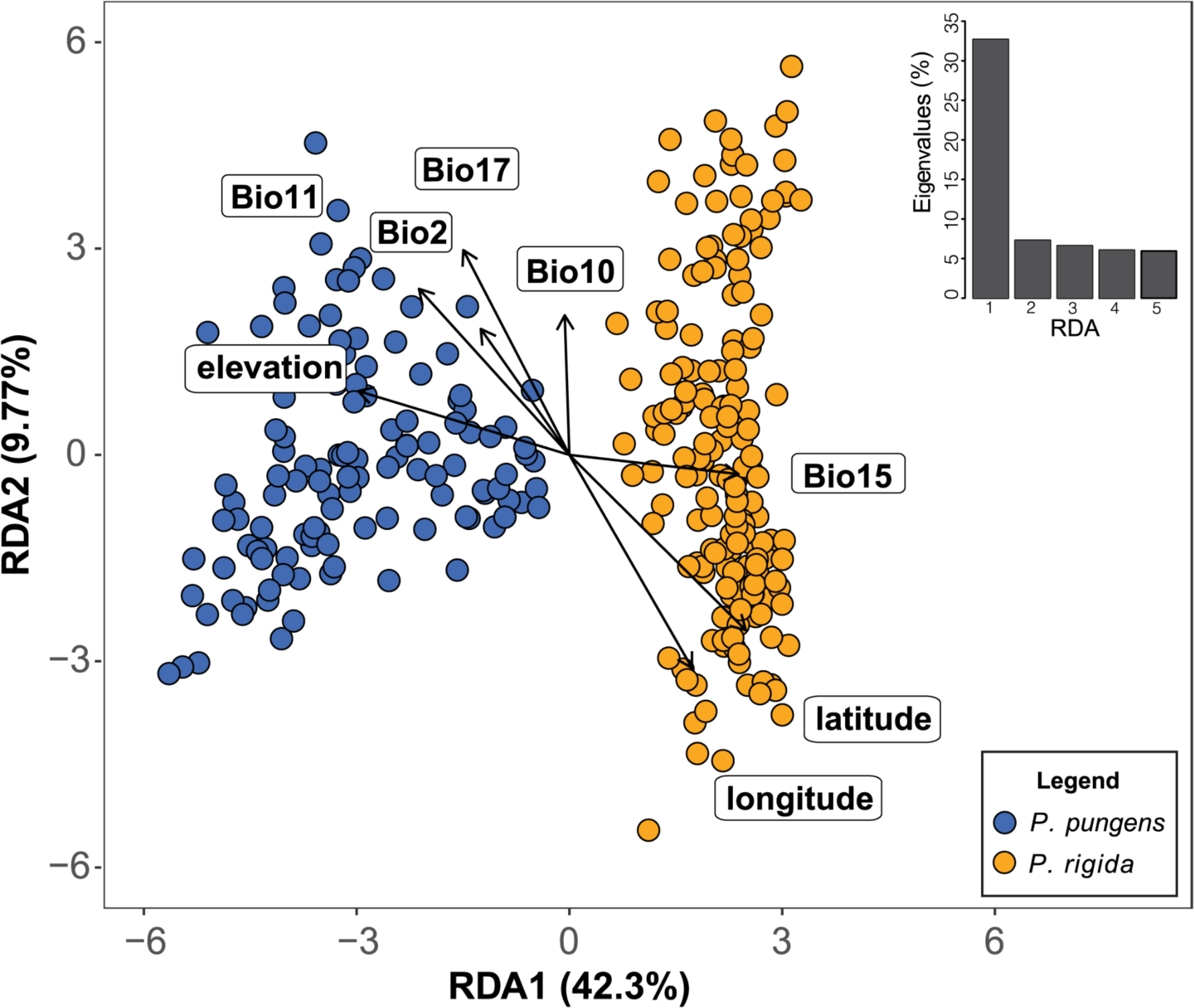
Redundancy analysis (RDA) of the multilocus genotypes for each tree with climate and geographic predictor variables (full model). Direction and length of arrows on each RDA plot correspond to the loadings of each variable.

Partitioning the effects of each predictor set revealed that climate independently (i.e., conditioned on geography) accounted for 31.93% of the explanatory variance. Geography independently (i.e., conditioned on climate) accounted for 34.10% of the explained variance. The confounded effect, due to the correlations inherent to the chosen geographic and climatic predictor variables, was 33.97%.

### Species distribution modeling

Because population structure within each of the focal species was not observed from our genetic data (i.e., no clear genetic clusters were identified), we produced SDMs using occurrence records across the full distributional range of each species. The best-fit SDM for *P. pungens* used a linear and quadratic feature class with a 1.0 regularization multiplier, while the SDM for *P. rigida* used a linear, quadratic, and hinge feature class with a regularization multiplier of 3.0. The AUC associated with the training data of the *P. pungens* and *P. rigida* SDMs was 0.929 and 0.912, respectively. Metadata, data inputs, outputs, and statistical results for model evaluation are available in Online Resource 2. The climatic variables with the highest permutation importance were mean temperature of the coldest quarter (Bio11) and precipitation seasonality (Bio15) which contributed 41.1% and 39.7% to the *P. pungens* SDM and 19.5% and 62.4% to the *P. rigida* SDM, respectively. Of the five climate variables included in the RDA, Bio15 and Bio11 had the highest loadings along RDA axis 1, helping to explain differences across species. The tandem reporting of Bio15 and Bio11 as important to both genetic differentiation and species distributions is indicative that these climatic variables contributed to the divergence of these two species.

Distributional overlap was observed in all analyzed SDMs at each of the four time points, therefore all four hypotheses stated that gene flow occurred between the LIG and present day (Fig. 4). Current SDMs indicated a larger area of suitable habitat for *P. rigida* (11,128 grid cells had > 0.5 habitat suitability) compared to *P. pungens* (6,632 grid cells) with 14.1% overlap in distributional predictions (Fig. 4). According to the SDM predictions, the areas of high habitat suitability shifted substantially over time for both species, with overlapping areas of suitable habitat exhibiting some of these fluctuations, as well. SDM ensembled predictions for HOL indicated the highest overlap (21.2% of grid cells with > 0.5 habitat suitability), while LGM ensembled predictions indicated the lowest overlap (9.1%). Likewise, calculations of overlap from full distributional predictions were the lowest (Schoener’s *D* = 0.217) for LGM followed by the LIG (Schoener’s *D* = 0.288). The highest distributional overlap was associated with the current SDM (Schoener’s *D* = 0.612; Fig. S5). Raster plots associated with the SDM predictions across the four time points (LGM and HOL ensemble predictions) and species are in Online Resource 4, Fig. S5.

**Fig. 4.**
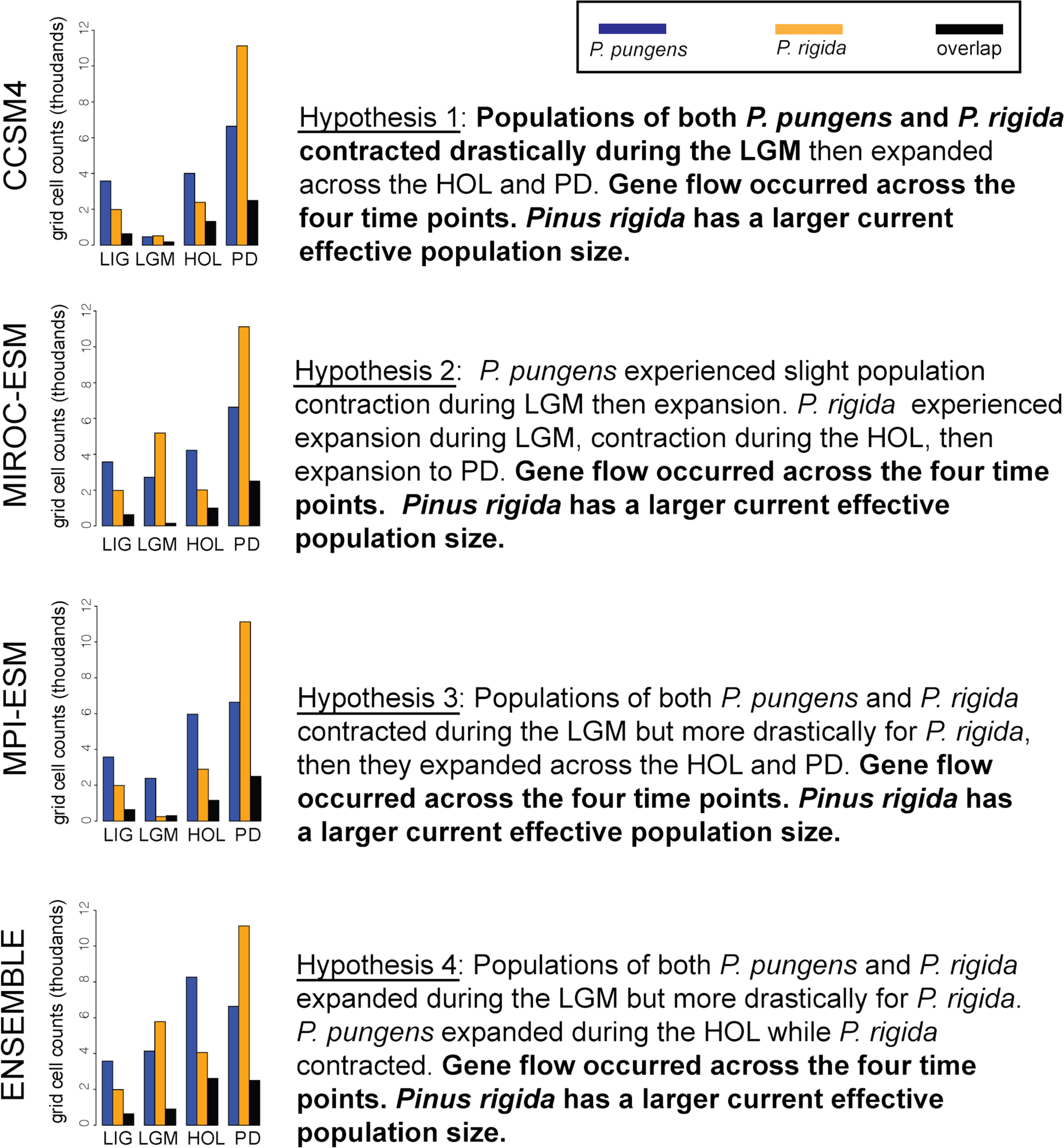
Hypotheses associated with each SDM - GCM model prediction versus the ensemble SDM prediction based on relative grid cell counts of high habitat suitability ( > 0.5) for *P. rigida*, *P. pungens*, and overlap across four time periods (LIG, LGM, HOL, and PD). Bolded text were statements supported by the best-fit model of demographic inference.

LGM predictions across the three GCMs varied substantially in terms of where and to what extent there was suitable habitat. We observed drastic reduction in suitable habitat for both species from predictions associated with the CCSM4 GCM. MPI-ESM associated predictions indicated reductions for *P. rigida*, while MIROC associated predictions indicated habitat expansion for *P. rigida* since the LIG. As found in Varela et al. (2015), the use of mean diurnal range (Bio2) and precipitation seasonality (Bio15) in historical SDM modeling for the LGM led to very different predictions across GCM types making averaged predictions (i.e., the ensemble approach) potentially misleading. We have provided model predictions associated with each LGM-GCM in Online Resource 4 (Fig. S6). Calculations of overlap from all LGM-GCM predictions (range = 2.0 - 18.3%) were lower than overlap estimates from other time periods providing some indication of consistency and usefulness to the widely-implemented ensemble technique. For the HOL, predictions were more similar across GCMs with overlap varying between 13.1 and 20.5% (Online Resource 4, Fig. S7). Hypotheses associated with each GCM and the ensemble are presented in Fig. 4.

The ensembled prediction for *P. pungens* and *P. rigida* during the LGM shows multiple potential refugial areas that overlap (Fig. S5). From the MIROC-ESM GCM-based model predictions, interspecific gene flow during the LGM may have been possible just south of the glacial extent, but CCSM4 and MPI-ESM GCM-based predictions (Fig. S6) indicate two, small overlapping refugial regions farther south than where either species currently occurs. Ensembled distributions for *P. pungens* and *P. rigida* during the HOL were proximal to each other, with high habitat suitability west of and along the Appalachian Mountains (Fig. S5). These distributions may have promoted both intraspecific and interspecific gene flow to occur ∼6 kya.

### Demographic modeling

The best replicate run (highest composite log-likelihood) for each of the thirteen modeled divergence scenarios, their associated parameter outputs, and ΔAIC (AICmodel i – AICbest model) are summarized in Online Resource 3. A model that allowed changes in both effective population size and rate of symmetrical gene flow across two time periods (PSCMIGCs) best fit the 2168 SNP data set (Table 2) and had small, normally distributed residuals (Fig. S8). This model was 20.84 AIC units better than the second best-fit model (PSCMIGs; Table 3), which inferred change in population size estimates across two time intervals but inferred only one, constant symmetrical gene flow parameter across time intervals.

**Table 3.** Results of model fitting for thirteen representative demographic models of divergence. Models are ranked by the number of parameters (*k*). Log-likelihood (log*L*) and Akaike information criterion (AIC) are provided for each model. Model details are given in the footnote.

| Model | $k$ | $\log L$ | AIC |
| --- | --- | --- | --- |
| SI | 3 | -2254.18 | 4,514.37 |
| MIGs | 4 | -2201.51 | 4,411.02 |
| MIGa | 5 | -2210.81 | 4,431.62 |
| SCs | 5 | -2213.93 | 4,437.86 |
| SGFs | 5 | -2229.65 | 4,469.30 |
| SCa | 6 | -2238.03 | 4,488.06 |
| SGFa | 6 | -2241.07 | 4,494.14 |
| PSC | 6 | -2277.78 | 4,567.56 |
| PSCSCs | 7 | -2178.16 | 4,370.32 |
| PSCMIGs | 7 | -1866.42 | 3,746.84 |
| **PSCMIGCs** | **8** | **-1853.99** | **3,726.00** |
| PSCMIGa | 10 | -2117.91 | 4,251.82 |
| PSCMIGCs_T3 | 12 | -1899.56 | 3,823.12 |

SI, strict isolation; MIGs, symmetrical gene flow; MIGa, asymmetrical gene flow; SCs, secondary contact with symmetrical gene flow; SCa, secondary contact with asymmetrical gene flow; SGFa, speciation with asymmetrical gene flow; SGFs, speciation with symmetrical gene flow; PSC, population size change; MIGCs, change in rate of symmetrical gene flow; T3, for three time intervals. The best-fit model is in bold.

Initial divergence was estimated to be 2.74 mya (95% CI: 2.25 – 3.24). The first time interval during divergence (*T1*) lasted 98.7% of the total divergence time with symmetrical gene flow (*Mi*) occurring at a rate of 48.6 (95% CI: 33.1 – 64.1) migrants per generation (Fig. 5). The effective size of the ancestral population (*Nref*) was 36,137 (95% CI: 31,367 – 40,908; Fig. 5) prior to divergence. For most of the divergence history, *P. pungens* had an effective population size of *NP1* = 1,024,573 (95% CI: 140,601 - 1,908,546) while *P. rigida* had a relatively smaller, but still large, effective size of *NR1* = 758,920 (95% CI: 214,423 - 1,303,417). The second time interval (*T2*) during divergence was estimated to have begun 35.2 kya (95% CI: 32.9 - 37.4) when effective population sizes decreased instantaneously to 3,448 (95% CI: 3,226 - 3,669) for *P. pungens* (*NP2*) and 3,935 (95% CI: 3,679 - 4,191) for *P. rigida* (*NR2*). During this time interval, the relative rate of symmetrical gene flow dropped from 48.6 to 38.4 (95% CI: 35.7 – 41.1) migrants per generation.

**Fig. 5.**
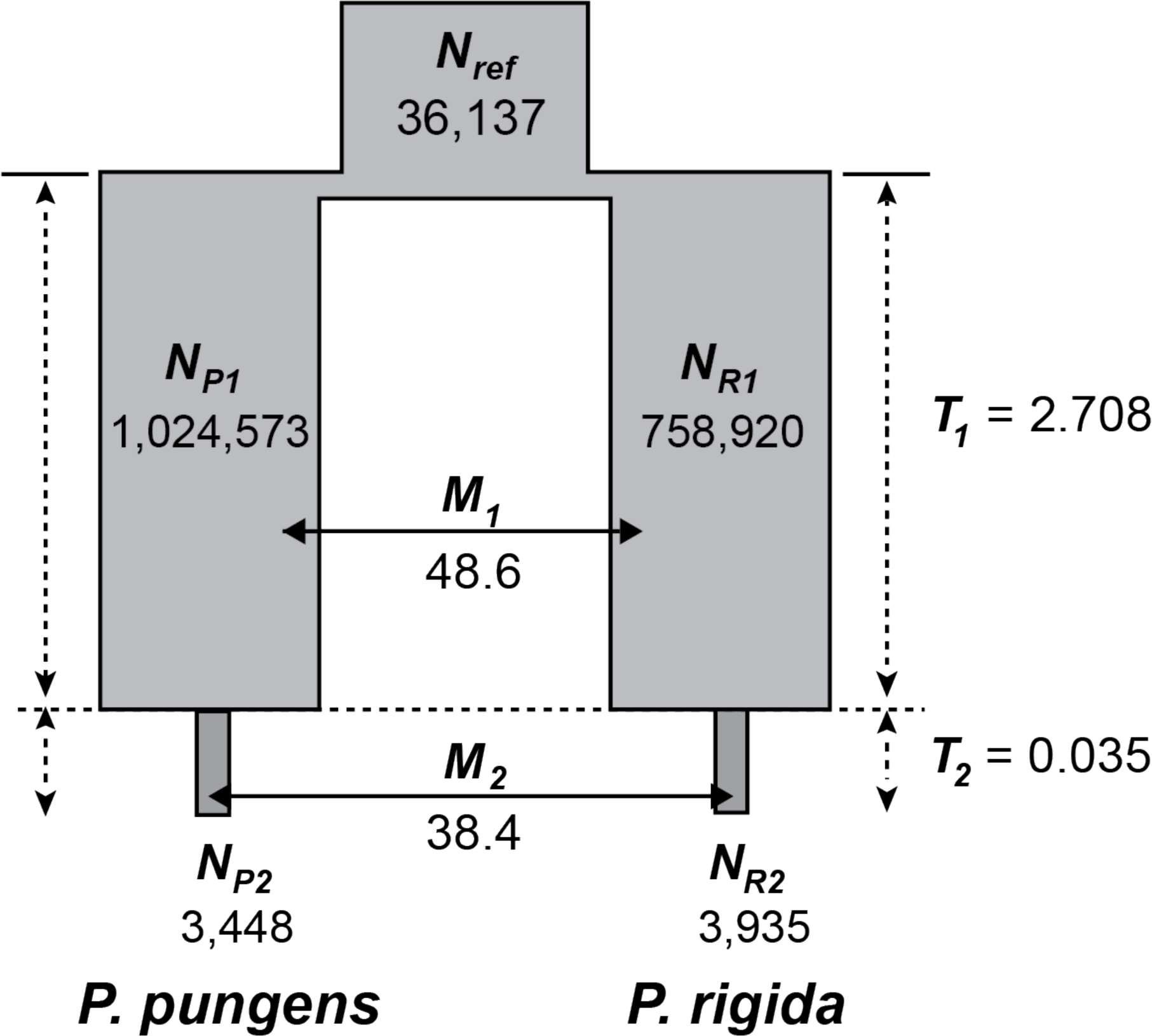
The best-fit model (PSCMIGCs) and unscaled parameter estimates from *δαδi* analysis. Time intervals (*Ti*) are represented in millions of years and associated with lineage population sizes (*Ni*) and a specific rate of symmetrical gene flow (*Mi*).

## Discussion

Using a multidisciplinary approach, we demonstrated that the divergence history of *P. pungens* and *P. rigida* involved a complex mixture of population size changes linked to changing climates, as well as changing rates of gene flow as quantified using the number of migrants per generation. We also demonstrated that consideration of each GCM-based SDM prediction is important to hypothesis formation for phylogeographic and demographic inference studies as the more widely employed method of ensembling historical SDM predictions can be misleading, especially when inferences include population size change. All four of our SDM hypotheses were supported in terms of gene flow occurrence since the LIG, but only the SDM hypothesis for population size change since the LIG (CCSM4, Hypothesis 1 in Fig. 4) was supported by genetic data. The best-fit demographic model using 2168 SNPs as summarized using the multidimensional site frequency spectrum indicated initial divergence to have occurred 2.74 mya, an estimate similar to the one inferred in Saladin et. al. (2017; 2.66 mya). Our best-fit model also indicated a large reduction in effective population size which coincided with a reduction in gene flow during the last glacial period (∼10,000 years before the last glacial maxima). A three-epoch model to test SDM observations of expansion since the LGM was included, but model fit did not improve. This could be due to the more pronounced impact of a recent bottleneck to site frequency spectrum patterns or that our data simply did not capture expansion from inferred bottlenecks.

### Climate drives divergence

The total divergence time inferred for *P. pungens* and *P. rigida* (2.74 mya) aligns with the onset of the Quaternary Period (∼2.6 mya), a time period widely recognized as driving adaptations to seasonality for many temperate species (Dobzhansky 1950; Savolainen et al. 2004; Jump and Penuelas 2005; Williams and Jackson 2007; Bonebrake and Mastrandea 2010). For *P. pungens* and *P. rigida*, precipitation seasonality (Bio15) was important to genetic differentiation (RDA) and species distributions (SDMs) which strongly implies adaptations to seasonality were drivers of divergence. While SDM predictions are often scrutinized, few features (e.g., linear and quadratic) were used in the predictions associated with our best-fit SDMs. This suggests high model accuracy and thus dependable identification of climatic variables (e.g., precipitation seasonality) important to habitat suitability. Phenological traits have been linked to seasonal variation within various plant species of North America (Jump and Penuelas 2005), and differences in seasonality requirements for *P. pungens* and *P. rigida* (i.e., mean of 11.3 versus 14.2, respectively) likely explain the observed trait differences in seed size, reproductive age, timing of pollen release, and rates of seedling establishment across these two species (Zobel 1969; Della-Bianca 1990; Ledig et al. 2015).

Using niche and trait data, the phylogenetic inference of Jin et al. (2021) also identified precipitation seasonality (Bio15) as a driver of diversification in eastern North American pines along with annual mean temperature (Bio1), mean temperature of the wettest quarter (Bio8), elevation, and soil silt content. Although three of these variables were not included in our RDA, the two that were (i.e., precipitation seasonality and elevation) were most important to explaining species level genetic differences. In terms of distributional differences between these two species, narrow niche requirements for precipitation seasonality and elevation help explain the patchy distribution of *P. pungens* along the southern Appalachian Mountains, while contrastingly, populations of *P. rigida* may have evolved a response to increased precipitation seasonality during the Quaternary period. In a study of pinyon pine diversification, Ortiz-Medrano et al. (2016) suggested the response to seasonality as potentially linked to the evolution of plasticity. This could explain *P. rigida*’s less stringent niche requirements for precipitation seasonality and elevation, larger geographic distribution, greater trait variation, and proposed latitudinal expansion into northeastern North America (Ledig et al. 2015).

The evolution of fire-related traits in pines has been linked to the mid-Miocene period, but fire intensity and frequency in certain geographic regions have been cyclical in nature thus allowing the evolution of adaptive traits related to fire endurance, tolerance, or avoidance to be possible across multiple geologic time scales (e.g., He et al. 2012; Lafon et al. 2017; Jin et al. 2021). Fine-scale geographical distributions of our focal species are locally divergent across slope aspects in the Appalachian Mountains, with *P. pungens* primarily distributed on southwestern slopes and *P. rigida* primarily distributed on southeastern slopes (Zobel 1969). Currently, there is higher fire frequency and intensity on western slopes. The high levels of cone serotiny and fast seedling development associated with *P. pungens* are evolved strategies that confer population persistence in more active fire regimes (Zobel 1969). Although some northern *P. rigida* populations exhibit serotiny, the populations found along the southern Appalachian Mountains, and proximal to *P. pungens,* have nonserotinous cones and other traits consistent with enduring fire (e.g., thick bark and epicormics; Zobel 1969) as opposed to relying on it (Jin et al. 2021). With these factors in mind and the correlative evidence between fire intensity and level of serotiny presented across populations of other pine species (*P. halepensis* and *P. pinaster*; Hernandez-Serrano et al. 2013), we suspect genomic regions involved in the complex, polygenic trait of serotiny (Parchman et al. 2012; Budde et al. 2014) may have also contributed to the rapid development of reproductive isolation, likely postzygotically, between our focal species.

### Reproductive isolation can evolve rapidly during speciation

While *P. pungens* and *P. rigida* can be found on the same mountain and even established within a few meters of each other, mountains are heterogeneous, complex landscapes offering opportunity for niche evolution along multiple axes of biotic and abiotic influence for parental species and hybrids alike. The distances to disperse into novel environments are relatively short in these heterogeneous landscapes thus suggesting diversification could be more rapid as environmental complexity increases (Bolte and Eckert 2020). Mountains have rain shadow regions characterized by drought and thus more active fire regimes (Parisien and Moritz 2009). A host of adaptive traits in trees are associated with fire frequency and intensity (Pausas and Schwilk 2012). Among those, the genetic basis of serotiny is characterized as being polygenic with large effect loci in *P. contorta* Dougl. (Parchman et al. 2012) and in *P. pinaster* Aiton (Budde et al. 2014). Such genetic architectures, even in complex demographic histories such as the one described here, can evolve relatively rapidly to produce adaptive responses to shifting optima (e.g., Stetter et al. 2018; reviewed for forest trees by Lind et al. 2018), so that it is not unreasonable to expect divergence in fitness-related traits such as serotiny to also contribute to niche divergence and reproductive isolation. Considering large effect loci associated with serotiny were also associated with either water stress response, winter temperature, cell differentiation, or root, shoot, and flower development (Budde et al. 2014), serotiny may be a trait that contributes to widely distributed genomic islands of divergence thus explaining the development of ecologically based reproductive isolation between *P. pungens* and *P. rigida* amid recurring gene flow (Nosil and Feder 2012). Given that our focal species are reciprocally crossable to yield viable offspring (Critchfield 1963), it is likely that postzygotic ecological processes, such as selection for divergent fire-related and climatic niches, limits hybrid viability in natural stands as a form of reinforcement layered on the aforementioned prezygotic divergence of phenological schedules. Indeed, hybrids are rarely identified in sympatric stands (Zobel 1969; Brown 2021). Thus, it appears that niche divergence is associated with divergence in reproductive phenologies during speciation for our focal taxa. Whether niche divergence reinforces reproductive isolation based on pollen release timing or divergent pollen release timing is an outcome of niche divergence itself, however, remains an open question.

The rate of gene flow in our best-fit demographic model was reduced by approximately 10 migrants per generation providing evidence that prezygotic reproductive isolation may have strengthened during the glacial period. This reduction reflects a scenario of reduced effective population sizes, reduced rates of gene flow (*m*), or both. The rate of gene flow associated with a given time interval should also not be interpreted as constant. For example, Sousa et al. (2011) found that posterior distributions for the timing of gene flow parameters in demographic inference were highly variable across the simulations they performed making pulses of gene flow (i.e., a gene flow event occurring within a time frame of no active gene flow), as probable as constant, ongoing gene flow. This likely explains the high levels of gene flow inferred using *δαδi* with the empirical lack of frequent and identifiable hybrids in extant samples of each species (Fig. 2; Brown 2021). While acknowledging this blurs interpretation of parameter estimates for gene flow, a history with recurring gene flow events fits the narrative of prezygotic isolation being labile especially when geographical distributions or reproductive phenology are the factors involved. Indeed, observations of hybridization occurring between once prezygotically isolated species have been made and suggests phenological barriers such as timing of pollen release and flowering may not be permanently established and can shift towards synchrony in warming climates (Vallejo-Marín and Hiscock 2016).

### Climate instability reduces genetic diversity

Conifers typically have high levels of genetic diversity and low levels of population differentiation because of outcrossing, wind-dispersion, and introgression (Petit and Hampe 2006). *Pinus pungens* and *P. rigida* both have modest levels of genetic diversity within and across the populations we sampled, and no detectable within-species population structure given our genome-wide data. Our best-fit model inferred a drastic effective population size reduction (*P. pungens,* ∼99.7%; *P. rigida*, ∼99.5%) to have occurred 35 kya. Since then, climate has continued to oscillate between extreme warming and cooling events (Jackson & Overpeck 2000) and for geologic time intervals too short for species with long generation times and low migratory potential to sufficiently track causing a mismatch between the breadth of a species’ climatic niche and where populations are established (Svenning et al. 2015). This dynamic affects population persistence, reduces genetic variation within populations due to excessive mortality, and thus to some degree limits the potential for local adaptation in climatically unstable regions. The lack of IBD and IBE across the populations of our focal species can be explained in one of two ways, the mismatch described in Svenning et al. (2015) or the primarily nongenic regions investigated in our RADseq data reflect little to no structure. Our SDM predictions showed substantial shifts in habitat suitability since the LIG, providing evidence of high climate instability in temperate eastern North America during the Quaternary period. We acknowledge though that niche conservatism is an underlying assumption in historical SDMs, so our interpretations were done cautiously. Gene flow and local adaptation affect niche dynamics in various ways (Pearman et al. 2008), but neither of these processes were able to be accounted for in our SDMs, especially when intersecting them with knowledge from our genetic data.

From a theoretical standpoint, we anticipated the patchy, mountain top distribution of *P. pungens* to be characterized by strong patterns of population differentiation. Lack of structure in *P. pungens* could be attributed to long distance dispersal or a recent move up in elevation with genomes still housing elements of historical panmixia. Indeed, suitable habitat predictions during the HOL, just 6000 years ago, were rather contiguously distributed (Fig. S5 and Fig. S7) and may have allowed an increase in intraspecific gene flow. For *P. rigida* some structure differentiating the northern populations from those along the southern Appalachian Mountains was expected from an empirical standpoint because previously reported trait values in a common garden study led to identification of three latitudinally arranged genetic groupings (Ledig et al. 2015). Although structure analysis did not support groupings within *P. rigida*, our estimates for isolation-by-distance (IBD) yielded a correlation of 0.177 (*p* = 0.055) which is suggestive of structure. While this shows some differentiation across its distribution, pairwise *F*ST values were small and on average smaller than those between populations of *P. pungens* suggesting higher population connectivity in *P. rigida*. The three GCM-based SDM predictions for both *P. pungens* and *P. rigida* differed substantially but did consistently show two or three disjunct refugia where gene flow dynamics intraspecifically and interspecifically may have been affected. Even though genetic differences may have accumulated in these separate refugia, the SDM predictions for the HOL were more compact and contiguous for our focal taxa, providing greater potential for intraspecific gene flow across diverged populations and the reestablishment of interspecific gene flow under a warming climate.

## Future work and conclusions

The divergence history of *P. pungens* and *P. rigida* involved a complex interplay of recurring interspecific gene flow and dramatic population size reductions associated with changes in climate. Future detailed examinations of hybridization between *P. pungens* and *P. rigida* are needed to elucidate the role hybridization plays in the maintenance of species boundaries. Ideally, future research involving these two species would use a method that sufficiently captures genic regions and thus the genomic islands of divergence that are often associated with ecological speciation (Nosil and Feder 2012). It may also be of interest to conduct population genetic analyses from chloroplast and mitochondrial DNA to obtain resolved inferences of gene flow directionality (i.e., asymmetry) and population connectivity.

While more time, effort, and genomic resources are needed for us to accurately predict gains and losses in biodiversity or describe the development of reproductive isolation in conifer speciation, we must recognize that some montane conifer species will be disproportionately affected by future climate projections (Aitken et al. 2008) and time is of the essence in terms of capturing and understanding current levels of biodiversity. High elevational species such as *P. pungens* may already be experiencing a tipping point, but because *P. pungens* is a charismatic Appalachian tree with populations already threatened by fire suppression practices over the last century, conservation efforts have begun through seed banking (Jetton et al. 2015) and prescribed burning experiments of natural stands (Welch and Waldrop 2001). Our contributions to these conservation efforts include genome-wide population diversity estimates for *P. pungens* and *P. rigida* and a demographic inference scenario that involves a long history of interspecific gene flow. In conifer species of the family *Pinaceae*, there are multiple accounts of introgression occurring through hybrid zones (De La Torre et al. 2014; Hamilton et al. 2015; Menon et al. 2018). The implications of introgression are far-reaching, as it leads to greater genetic diversity and thus a greater capacity for adaptive evolution. Trees are often foundation species in many plant communities, so understanding a population’s potential to withstand environmental changes provides some insight into the future stability of the ecological communities dominated by these charismatic plant taxa.

## Author contributions

CB performed field sampling of *Pinus rigida,* data analyses, and modeling. TF processed the genetic data and advised statistical analyses. CF led the field sampling and library prep for *Pinus pungens*, and AJE assisted with data analyses. All authors contributed to the writing of this manuscript.

## Data Archiving Statement

Raw reads generated during this study are available at NCBI SRA database under BioProject: PRJNA803632 (Sample IDs: SAMN25684544 – SAMN25684843). Python scripts for demographic modeling and R scripts for genetic analyses and producing SDMs are available at www.github.com/boltece/Speciation_2pines.

## Supplemental material

Online Resource 1: Summary statistics per SNP, sampled tree, and population Online Resource 2: Metadata and files needed to reproduce SDMs

Online Resource 3: Results from demographic inference and parameter unscaling Online Resource 4: Supplemental figures

## Statements and Declarations

The authors have no financial or proprietary interests in any material discussed in this article. Field sampling permits at National Parks and State Parks were obtained prior to collecting leaf tissue: ACAD-2017-SCI-007, BLRI-2013-SCI-0027, GRSM-2017-SCI- 2028, ZZ-RCP-112514, and SFRA-1725.

## Supporting information

Online Resource 1

Online Resource 2

Online Resource 3

Online Resource 4

## Acknowledgements

This research was funded by Virginia Commonwealth University (VCU) Department of Biology, VCU Integrative Life Sciences, and National Science Foundation (NSF) awards to Andrew J. Eckert (NSF-EF-1442486) and Christopher J. Friedline (NSF-NPGI-PRFB- 1306622). We thank Mitra Menon and Rebecca Piri for their assistance with field sampling, Brandon Lind for providing computational support, and the undergraduate researchers who helped with the DNA extraction protocol: Kaylyn Carver, Casey Harless, and Leslie Ranson. We also thank the VCU Center for High Performance Computing for providing computational resources with which we made demographic inferences.

