## Supplementary material for "Divergence amid recurring gene flow: complex demographic histories for two North American pines (*Pinus pungens* and *P. rigida*) fit growing expectations among forest trees": Online Resource 4

Journal: TGG	Year: 2022

This resource includes supplemental figures mentioned in the main text

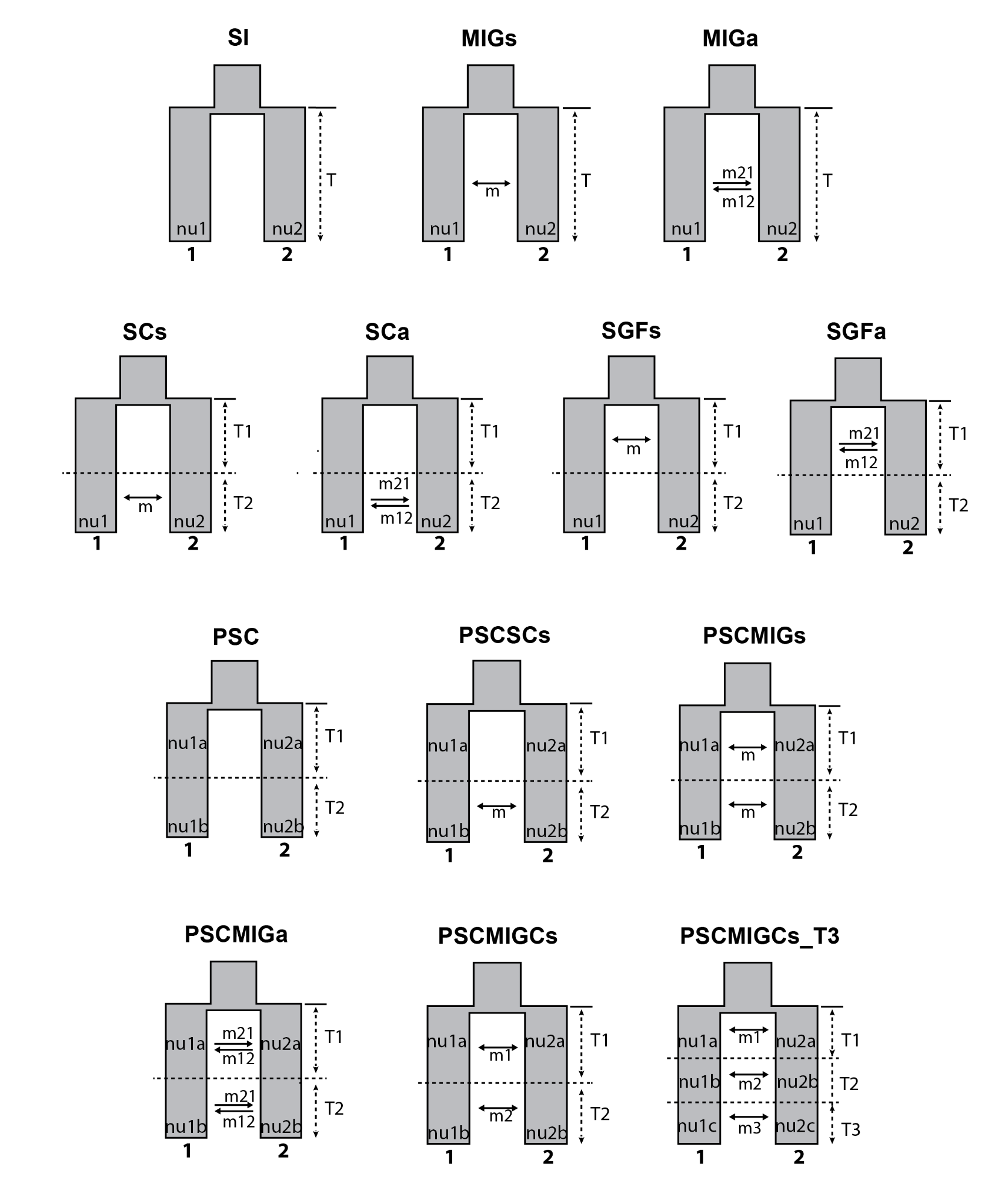

**Fig. S1** The thirteen divergence scenarios tested within the program $\partial\alpha\partial i$. Parameter estimates for the best run (highest log likelihood) of each scenario (model) are summarized in Online Resource 3

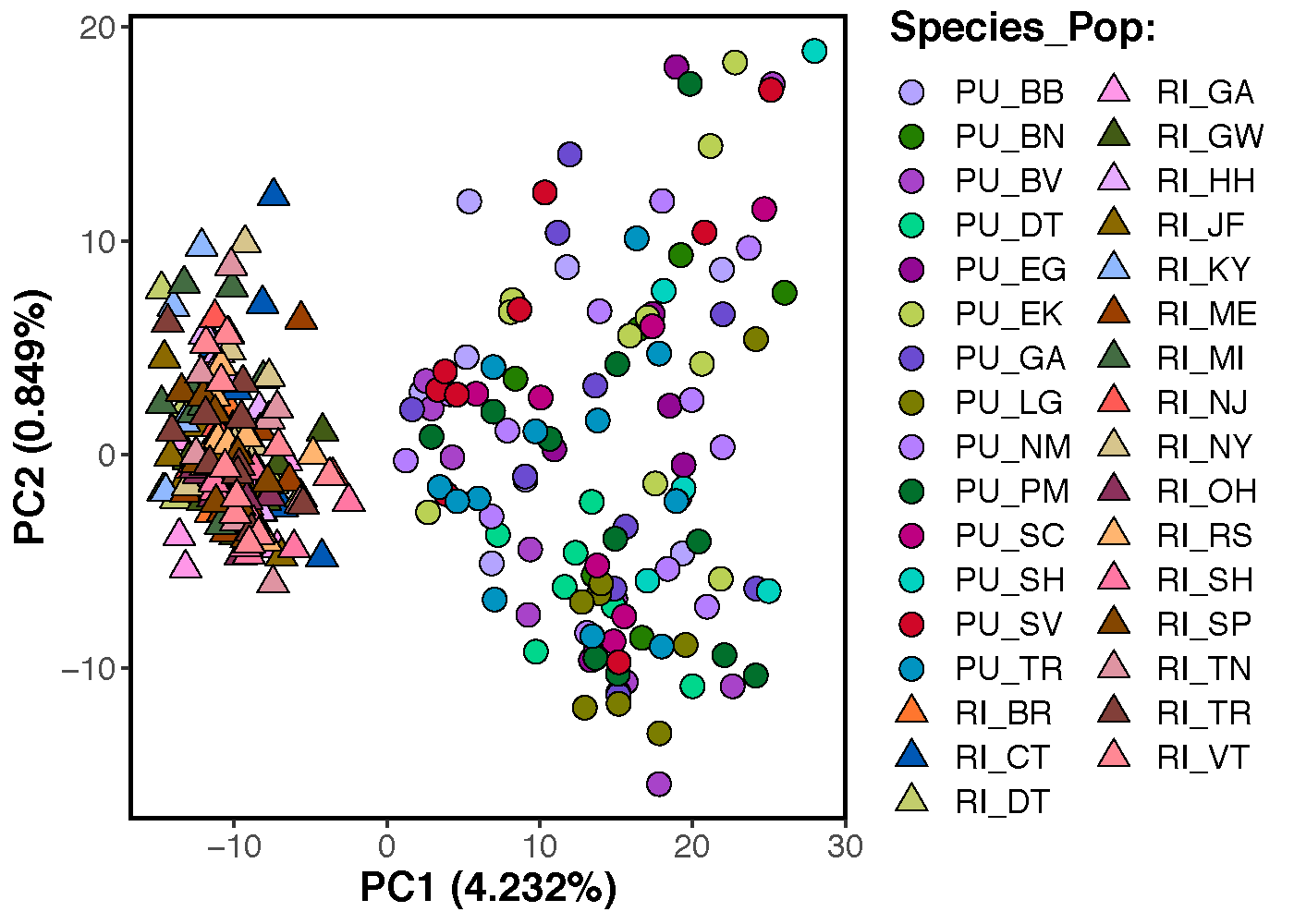

**Fig. S2** Principal component analysis (PCA) of 300 *P. rigida* and *P. pungens* trees labeled by population assignment

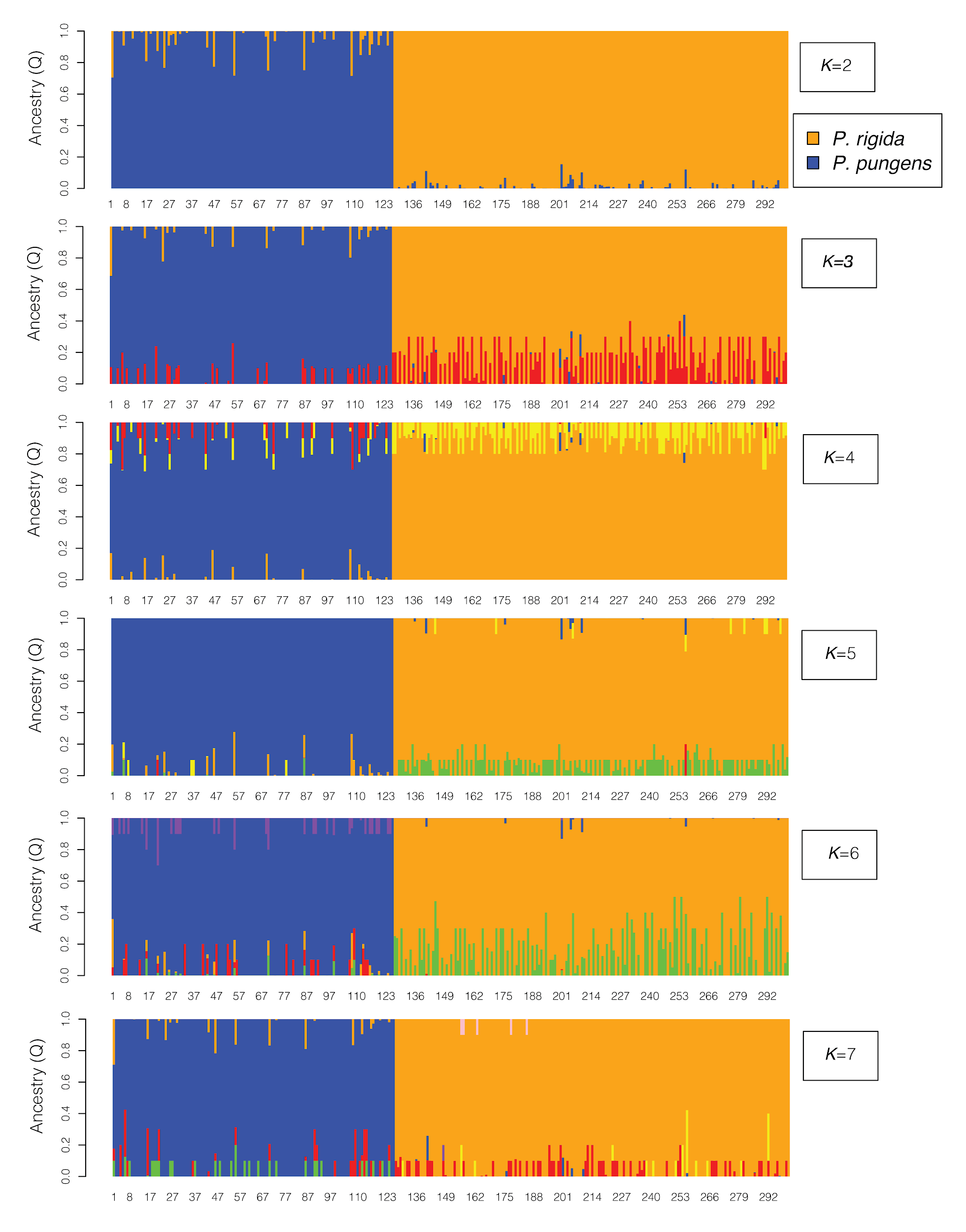

**Fig. S3** Individual based assignments of admixture from analysis of *fastSTRUCTURE* for K =2 through K =7. The plot associated with each value of K represents the averages assignments for each individual across 10 replicate runs

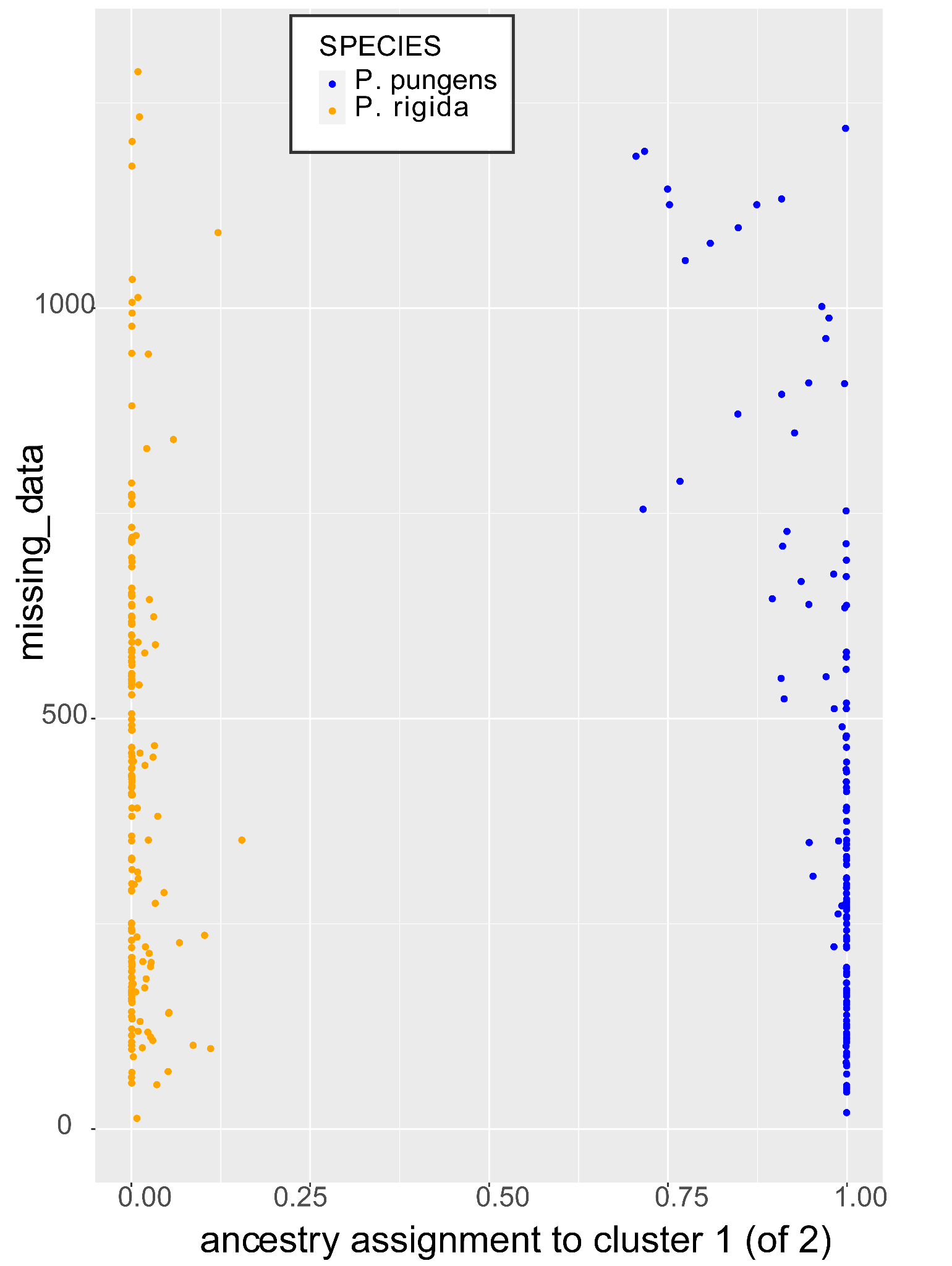

**Fig. S4** Distribution of missing data across the sampled trees in relation to ancestral coefficients (from K = 2). Blue circles to the right are samples of *P. pungens*. Orange circles to the left are samples of *P. rigida*

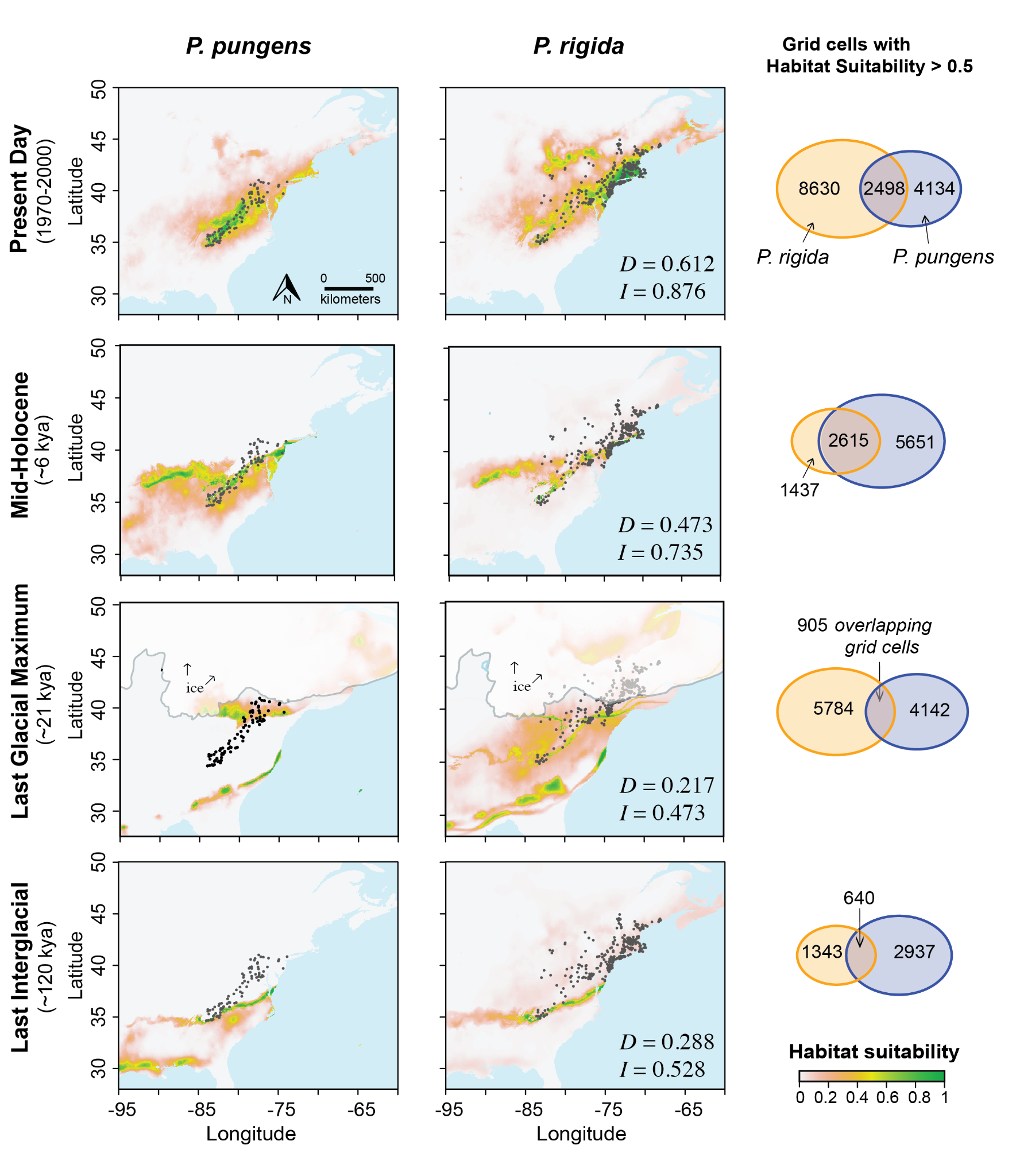

**Fig. S5** Species distribution model (SDM) predictions across four time points for *P. pungens* and *P. rigida*. Measures of raster overlap in terms of Schoener’s *D* and Warren’s *I* index between the models of each species, and at each time point, are presented in the bottom right corner of the prediction plots for *P. rigida.* Venn diagrams illustrate the number of grid cells with moderate to high habitat suitability scores (> 0.5) for each SDM at a given time point, as well as the number of shared, or overlapping, grid cells. Blue Venn diagram ovals show grid cell counts from the *P. pungens* SDM, and orange Venn diagram ovals show grid cell counts from the *P. rigida* SDM for the aligning time point (denoted on the left side). Habitat suitability distributions for LGM and HOL depict ensembled predictions. Glacial extent data (labeled ice in LGM plots) for 18 kya was provided by Dyke (2003)

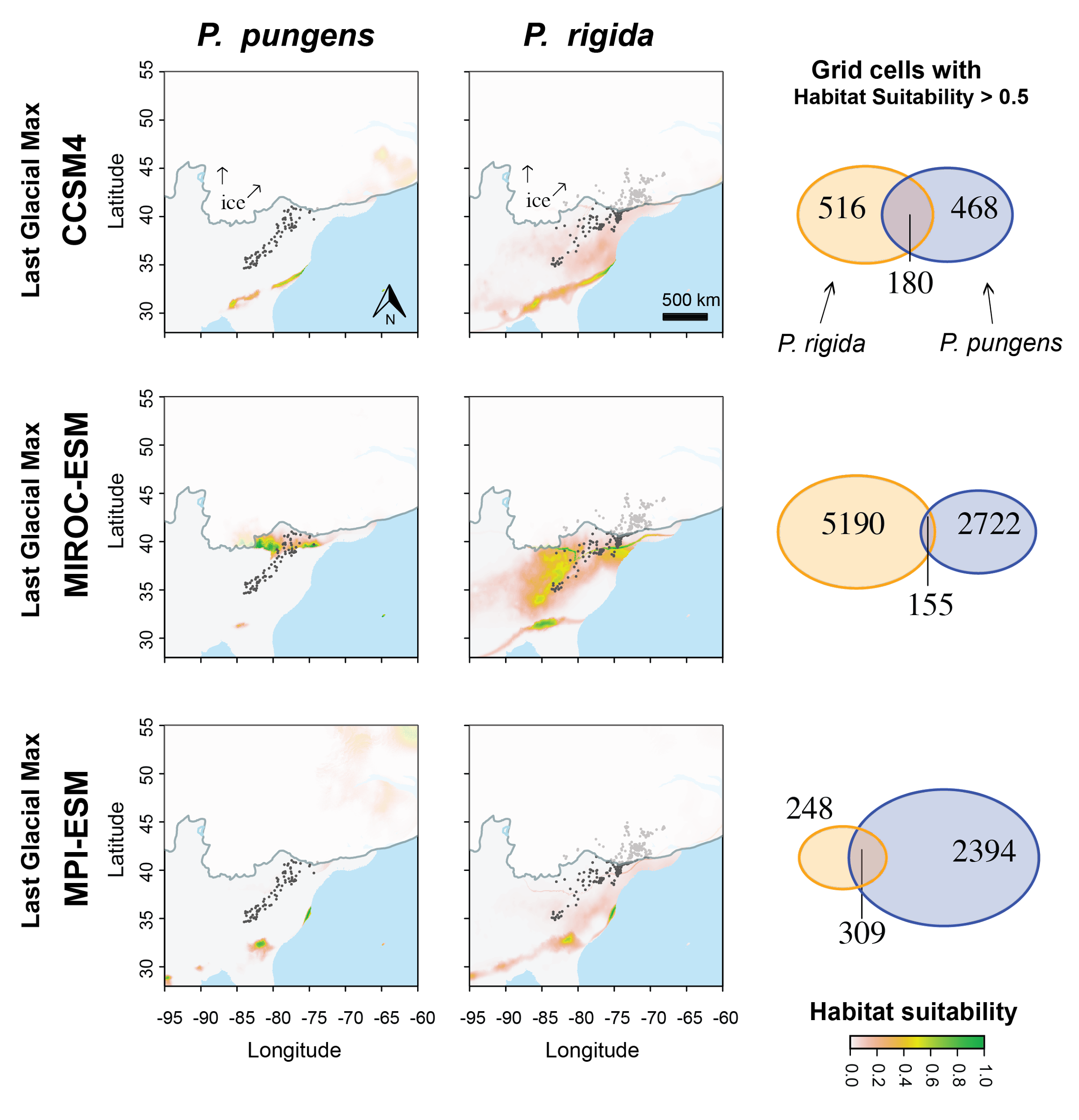

**Fig. S6** Last Glacial Maximum (LGM, ~21 kya) model predictions from each GCM (CCSM4, MIROC-ESM, and MPI-ESM) and Venn Diagram quantifications for size of suitable habitat (population size) and species overlap (gene flow).

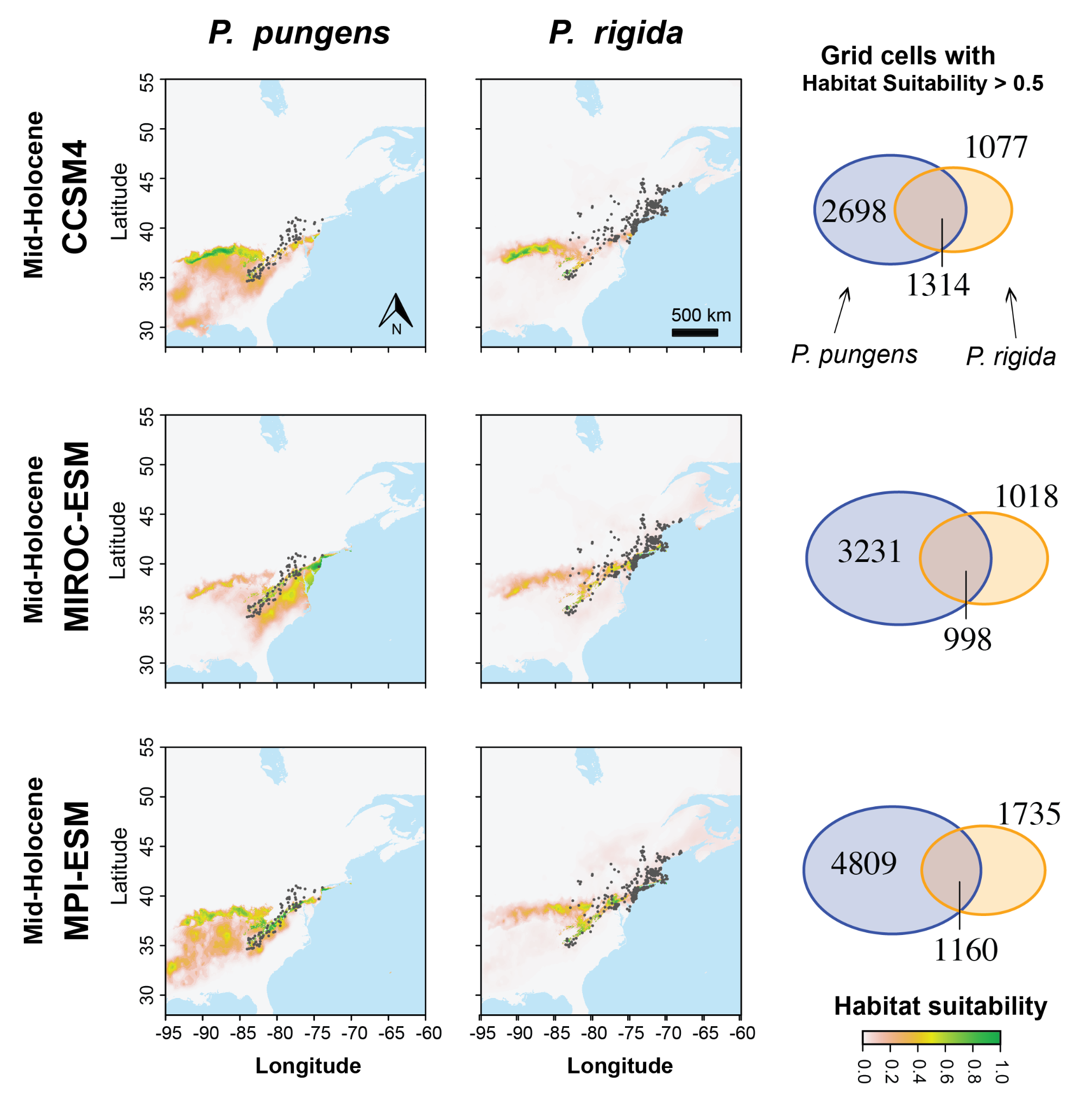

**Fig. S7** Mid-Holocene (~6 kya) model predictions from each GCM (CCSM4, MIROC-ESM, and MPI-ESM) and Venn Diagram quantifications for size of suitable habitat (population size) and species overlap (gene flow).

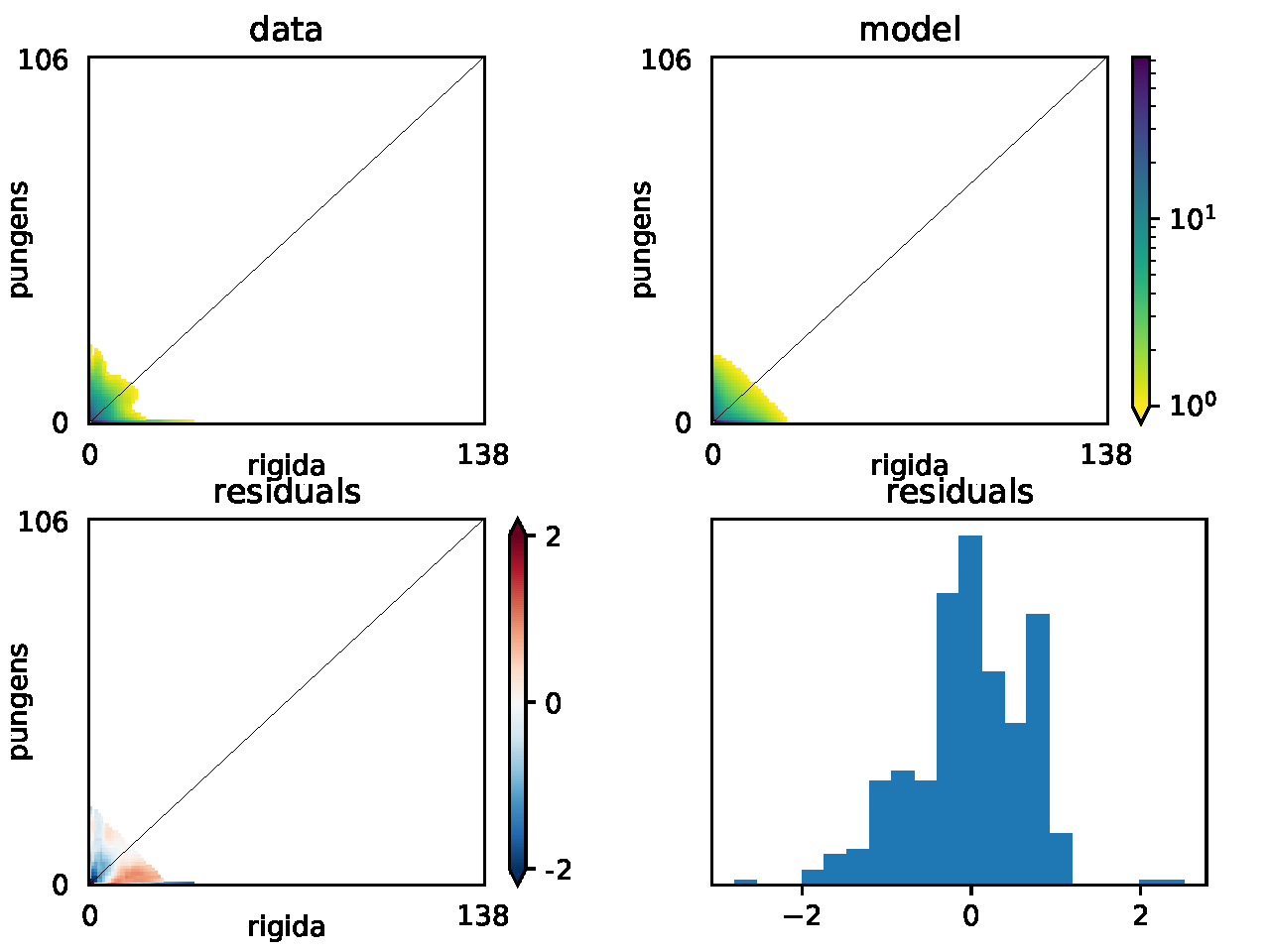

**Fig. S8** Presentation of data-model fit to the PSCMIGCs model run with highest log likelihood. Residuals between the data and model are shown in bottom row plots.
